# Linking the kinetic mechanism to structural dynamics required for nucleotide hydrolysis by an alphavirus nsP2 RNA helicase

**DOI:** 10.64898/2026.05.08.723793

**Authors:** Kacey M. Talbot, Yuan-Wei Norman Su, Jake B. Royster, David W. Gohara, Arash Firouzbakht, Marynna N. McLean, Bose Muthu Ramalingam, Timothy M. Willson, Jamie J. Arnold, Craig E. Cameron

## Abstract

RNA helicases encoded by positive-strand RNA viruses are essential for genome replication, yet the specific biological functions and mechanochemical basis underlying these functions remain poorly defined. Progress has been limited by the difficulty of resolving individual catalytic steps under single-turnover conditions, which are often experimentally inaccessible for viral enzymes. Alphaviruses replicate within membrane-bound spherules that may alter local metabolite concentrations, raising the possibility that the enzymatic properties of alphaviral proteins differ from those of viruses with greater cytosolic exposure. Here, we present a kinetic and binding analysis of full-length non-structural protein 2 (nsP2) from Chikungunya virus, a multifunctional superfamily 1B NTPase and RNA helicase. Purified nsP2 binds nucleoside triphosphates with high affinity, exhibiting equilibrium dissociation constants in the single digit micromolar range. This property enabled single-turnover, pre-steady-state, and isotope-trapping experiments that are rarely feasible for viral helicases. These analyses identified two sequential conformational-change steps required for nucleotide hydrolysis. Molecular dynamics simulations suggest tightening of the RecA1 and RecA2 domains upon ATP binding followed by compaction of the enzyme mediated by interactions between the 1B subdomain and RecA2 domain. Product inhibition patterns support random release of ADP and inorganic phosphate, with relative binding affinities indicating that ADP dissociates first. The reaction is irreversible. Although nsP2 binds RNA tightly, strand separation under single-turnover conditions is too slow to represent ATP-driven unwinding, instead likely reflecting formation of an unwinding-competent nsP2-RNA complex. Together, these findings establish a quantitative framework for nsP2 function and provide a roadmap for mechanistic studies of alphaviral helicases.

**Graphical Abstract:** 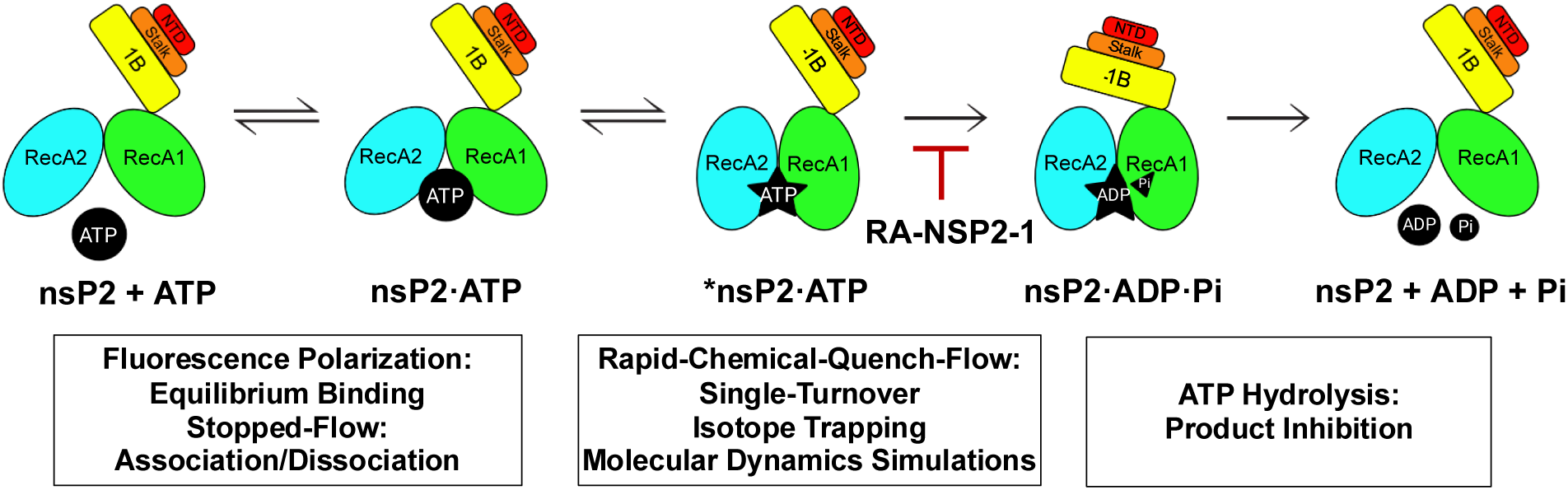

## Introduction

Most positive-strand RNA viruses of mammals encode an RNA helicase in their genomes that is essential for genome replication and virus multiplication (1,2). Although loss of helicase activity abrogates virus multiplication, the specific actions of the helicase required during the virus lifecycle are largely unknown (1,2). Generally, helicases equilibrate at the site for unwinding when bound to an RNA substrate (3,4). Subsequent nucleotide binding and hydrolysis, often reiteratively, leads to translocation of the helicase on RNA to promote unwinding of dsRNA, remodeling of RNA structures, displacement of proteins, among several other possibilities (5,6).

Despite extensive study, how the energy of nucleotide hydrolysis is coupled to unwinding has yet to be fully described for any helicase. One major limiting factor is the inability to assess both nucleotide hydrolysis and nucleic acid unwinding under single-turnover conditions, which are required to resolve individual chemical and conformational steps. For single-turnover conditions to be achieved, the concentration of enzyme must be on the order of 10-fold higher than the equilibrium dissociation constants (*K*_d_) for both the nucleotide and nucleic acid substrates. Achieving these conditions has been complicated by the low affinity (10-100 μM) of many helicases for their nucleoside triphosphate substrate(s), which would demand the use of enzyme concentrations in the 100-1000 μM range (4,7-9). Such enzyme concentrations are not easily realized experimentally.

Alphaviruses replicate their genomes in membrane structures termed spherules (10). These unique structures are formed by the assembly of non-structural proteins, particularly non-structural protein 1 (nsP1) on cellular membranes (2). For Chikungunya virus (CHIKV), spherules form on the plasma membrane (10). The viral replication machinery, including the RNA helicase, localizes to the “n k” of these spherules, where genome replication and associated RNA processing reactions occur (10). While the molecular details of the macromolecular assembly forming spherules are emerging, gaps remain in our understanding of the organization and composition of these structures (2,10,11). However, the current consensus opinion is that the interior of the spherule is not in direct contact with the cytoplasm, which may preclude viral proteins and genome replication intermediates from being surveilled by host responses (2,10). Importantly, such compartmentalization may also alter the effective concentrations of biomolecules relative to the cytoplasm, including nucleotides, with direct implications for enzymatic mechanism and regulation.

CHIKV encodes a superfamily 1b (SF1B), 5’→3’ nucleoside triphosphate hydrolas (NTPase), RNA helicase, and RNA triphosphatase (RTPase) in the amino-terminal domain of its non-structural protein 2 (nsP2) (12). The central domain of nsP2 contains a cysteine protease required for polyprotein processing (12). The carboxy-terminal domain contains a methyltransferase-like domain with no reported enzymatic activity or clearly defined function (12). Although the only structural information available for full-length nsP2 derives from small-angle X-ray scattering (13), higher resolution crystal structures are available for the isolated helicase (14) and protease (15) domains. While the helicase domain exhibits NTPase activity, helicase activity has not been robustly observed in vitro (13,16). Moreover, quantitative kinetic, equilibrium, and/or thermodynamic analysis of the nsP2-encoded NTPase and/or helicase has not been performed for nsP2 from any alphavirus.

In this study, we expressed and purified full-length nsP2 from CHIKV. We used the purified protein to perform a comprehensive suite of substrate binding (equilibrium and kinetics) and enzyme kinetics (steady-state and pre-steady-state) studies. The equilibrium dissociation constant (*K*_d_) for nsP2 binding to nucleotide was in the single digit micromolar range, tighter than values reported for many other viral helicases. This high affinity permitted us to perform single-turnover and isotope-trapping experiments. These experiments allowed us to unambiguously assign a conformational change as the rate-limiting step for nucleotide hydrolysis. Nucleotide hydrolysis was irreversible. Patterns of adenosine diphosphate (ADP) and inorganic phosphate (P_i_) product inhibition were consistent with both products capable of binding to free enzyme competitively with respect to ATP, suggesting random release of products. However, the equilibrium dissociation constants for ADP and P_i_ were consistent with ADP dissociating first.

RNA also binds to nsP2 with high affinity. However, single-turnover unwinding experiments revealed a rate constant too slow to represent ATP hydrolysis-driven unwinding and likely represents formation of a productive, unwinding-competent nsP2-RNA complex. We discuss the implications of our kinetic analysis on the intracellular function and regulation of nsP2 within alphavirus replication compartments. Together, this study establishes a mechanistic framework for nsP2-encoded helicases and provides a roadmap for investigating nsP2 helicases from other alphaviruses, enabling elucidation of differences that may inform the biological role of this enzyme.

## Materials and Methods

### Materials

RNA oligonucleotides were from Horizon Discovery Ltd. (Dharmacon). T4 polynucleotide kinase was from ThermoFisher. [γ-^32^P] ATP (6,000 Ci/mmol), [α-^32^P] NTPs (ATP, CTP, GTP, and UTP), and [α-^32^P] dNTPs (dATP, dCTP, dGTP, and dTTP) (3,000 Ci/mmol), were from Revvity. Nucleoside 5’-triphosphates and 2’-deoxynucleoside 5’-triphosphates (ultrapure solutions) were from Cytiva. Mant-ATP and mant-ATPγS were from Jena Bioscience. All other reagents were of the highest grade available from MilliporeSigma, VWR, or Fisher Scientific.

### Construction of CHIKV nsP2 bacterial expression plasmids

The CHIKV nsP2 gene was codon optimized for expression in *E. coli* and obtained from Genscript. The amino acid sequence for nsP2 was derived from CHIKV (GenBank # CAJ90477.1). The gene was directly cloned into the pTWIN-STREP-6HIS-SUMO bacterial expression plasmid using BsaI and SalI (17). The final pTWST-STREP-6HIS-SUMO-CHIKV-nsP2 construct was confirmed by sequencing by Plasmidsaurus.

### Codon optimized sequence for CHIKV nsP2

5’ggaat at gaga a t g ggag at aaggtga g a aa a tga a gt gttggggaata ttggt cctttctcctcagactgtgttgcgtagccagaaattgtctttaatccatgctctggccgagcaagtaaaaacgtgtacgcataacggccgtgcaggccgttatgccgtagaagcgtatgacggacgtgtcttagtcccctccggatatgctatctcgccggaggactttcagagcttatcggagtcagccaccatggtgtacaatgagcgtgaattcgtaaatcgcaaattacaccacatcgcaatgcatggtcccgccctgaacaccgatgaagaatcctatgagttagtgcgcgccgaacgcacggagcatgagtatgtttatgacgtagaccaacgccgctgttgtaaaaaagaagaggcggcaggtttggttctggtaggagatcttacaaatcctccgtaccatgaattcgcatacgaggggttgaaaatccgtccagcctgcccctataaaattgctgttatcggggtgttcggtgttccaggctctgggaagagcgcgattattaagaatttagtaactcgccaagatcttgtcacttcaggcaaaaaggagaattgtcaagaaattaccacagatgtgatgcgtcagcgcggattggaaattagcgcgcgtactgtggatagcttgcttctgaacgggtgcaatcgccccgtggatgtcttatatgtggacgaggcgttcgcatgtcacagtggcaccctgttagcgctgattgccttggttcgtccacgtcagaaggtcgtattgtgtggcgatccaaaacagtgtgggtttttcaacatgatgcagatgaaagtaaactacaatcataatatctgtactcaggtttatcacaagtccatttcccgtcgctgtacgttgcccgtaacggccatcgtttcctcgttgcactacgaaggtaagatgcgcacgactaacgagtacaataagccaattgtagtagacacgactggcagcacaaaaccggaccctggcgatttagtattaacttgttttcgcggttgggttaagcaattgcagatcgactatcgcggatacgaggttatgacagcggccgctagtcagggattaactcgcaaaggggtatacgcggtacgccagaaggtcaatgagaatccattgtatgcctccacttctgaacacgtgaatgtcttgttaactcgtaccgagggcaaattagtttggaagaccctttcgggtgacccgtggattaaaacccttcagaacccacccaaggggaacttcaaggctacaatcaaggagtgggaagtggaacatgcatcgattatggcggggatttgctcacatcagatgacgtttgatacgtttcagaataaagccaacgtatgttgggccaaaagtcttgtgcccatcctggagactgctggcattaagcttaacgaccgtcagtggtcacaaattattcaggccttcaaagaggataaagcctactcccccgaggtagccttaaatgagatttgcacacgtatgtacggggtagatttggactctggccttttcagcaaaccattggtatcagtgtactatgccgataaccactgggacaaccgccctggggggaaaatgttcggattcaaccctgaggccgcaagtattctggaacgtaaatacccattcacgaaaggtaagtggaacattaacaagcagatctgcgttacgacacgccgtatcgaagatttcaaccctactactaatatcattccagcaaaccgccgtttgcctcactccttggtcgcagaacaccgcccagtaaaaggtgaacgcatggaatggttggtcaataaaattaatgggcatcatgttttgctggtgagcgggtacaatttagcacttccgacgaagcgtgtcactt gggttgcaccccttggggtccgcggggctgactatacttataatctggagcttggtttacctgcaacgttgggtcgttatgatcttgttgtcatcaatattcacactccctttcgcatccaccattatcaacaatgtgtcgatcatgctatgaaattgcagatgctggggggggattccttgcgcttattgaaacccggtggtagcctgttaattcgcgcatacggctatgccgaccgcacgtctgagcgcgttatctgcgttcttgggcgtaagttccgtagctcgcgtgcacttaaaccaccttgcgtgacgtcaaatacagagatgttctttctgtttagtaattttgataacggacgccgtaactttaccactcacgtaatgaacaaccaattgaatgctgcttttgtagggcaggtcactcgtgcaggctgt-3’

### Construction of nsP2 active site derivatives

Single-amino acid substitutions were introduced into the CHIKV nsP2 coding sequence of pTWIN-STREP-6HIS-SUMO-CHIKV-nsP2 bacterial expression plasmid using the Agilent QuikChange Site-Directed Mutag n sis Kit, following the manufacturer’s instru tions. Mutagenic primers were designed to introduce the desired codon substitutions while preserving *E. coli* codon optimization. The following single point mutations were constructed: K192L, K192R, S193A, D252A, D252N, E253A, and E253Q. All mutant plasmids were confirmed by full-length plasmid sequencing performed by Plasmidsaurus.

### Expression of CHIKV nsP2

*E. coli* BL21(DE3) competent cells were transformed with the pTWIN-STREP-6HIS-SUMO-CHIKV-nsP2 plasmids (WT and derivatives) for protein expression. BL21(DE3) cells containing the pTWIN-STREP-6HIS-SUMO-CHIKV-nsP2 plasmid were grown in 100 mL of media (NZCYM) supplemented with kanamycin (K25, 25 µg/mL) at 37 °C until an OD_600_ of 1.0 was reached. This culture was then used to inoculate 4 L of K75-supplemented ZYP-5052 auto-induction media at 37 °C. The cells were grown at 37 °C to an OD_600_ of 0.8 to 1.0, cooled to 15 °C and then grown for 36-44 h. After ∼40 h at 15 °C the OD_600_ reached ∼10 -15. Induction was verified by SDS-PAGE using TGX-stain free gels and visualized by using Bio-Rad ChemiDoc MP Imager. Cells were harvested by centrifugation (6000 x *g*, 10 minutes) and the cell pellet was washed once in 200 mL of TE buffer (10 mM Tris, 1 mM EDTA), centrifuged again, and the cell paste weighed. For large scale growths, 50 L growths were performed by the Auto-Induction method described above but using 50 L fermenters at the PSU CSL Behring Fermentation facility.

### Purification of CHIKV nsP2

Frozen cell pellets were thawed on ice and suspended in lysis buffer (50 mM sodium phosphate pH 7.8, 500 mM NaCl, 1 mM TCEP, 20% glycerol) with 5 mL of lysis buffer per 1 gram of cells. The cell suspension was lysed by passing through a Nano DeBEE Gen II Homogenizer using nozzle Z08 in reverse flow mode (BEE International) at 20,000 psi. After lysis, NP-40 was added to a final concentration of 0.1% (v/v). While stirring the lysate, polyethylenimine (PEI) was slowly added to a final concentration of 0.25% (v/v) to precipitate nucleic acids from cell extracts. The lysate was stirred for an additional 30 minutes at 4 °C after the last addition of PEI, and then centrifuged at 75,000 x g for 30 minutes at 4 °C. The clarified PEI supernatant was separated and to it solid ammonium sulfate was slowly added to 40% saturation while stirring. Granular ammonium sulfate was pulverized to a fine powder using a mortar and pestle before use. The solution was stirred for an additional 15-30 minutes at 4 °C after addition of ammonium sulfate. The ammonium sulfate precipitated material was pelleted by centrifugation for 30 minutes at 75,000 x *g* at 4 °C. The supernatant was decanted and the pellet was suspended in 50 mM sodium phosphate pH 7.8, 500 mM NaCl, 1 mM TCEP, 20% glycerol, and 10 mM imidazole. The resuspended ammonium sulfate pellet was then loaded onto a 5 mL Nuvia Ni-NTA column (Bio-Rad) at a flow rate of 1 mL/min (approximately 1 mL bed volume per 100 mg total protein) equilibrated with buffer A (25 mM HEPES pH 7.5, 500 mM NaCl, 20% glycerol, 1 mM TCEP) with 10 mM imidazole using a Bio-Rad NGC System. After loading, the column was washed with twenty column volumes of buffer A with 10 mM imidazole and 5 column volumes of buffer A with 50 mM imidazole. The protein was eluted with 5 column volumes of buffer A with 500 mM imidazole. Ulp1 (1 µg per mg) was added to the eluted protein to cleave the SUMO fusion and was dialyzed overnight against 1 L buffer B (25 mM HEPES pH 7.5, 1 mM TCEP, 20% glycerol containing 500 mM NaCl (membrane employed a 12 -14,000 Da MWCO). After dialysis the dialyzed sample was diluted in Buffer B to a final NaCl concentration of 80 mM. The sample was loaded onto a 5 mL UNOsphere Q column in tandem with a 5 mL UNOsphere S column (1 mL bed volume/25 mg of protein) at 1 mL/min. The protein passed through the Q column and bound to the S column. Both columns were washed with two column volumes of Buffer B containing 80 mM NaCl, the S column was subsequently washed with 10 column volumes of Buffer B containing 80 mM NaCl. The protein was eluted from the S column using a linear gradient (6 column volumes) from 80 to 1000 mM NaCl in Buffer B. Fractions containing nsP2 protein were pooled based on purity from SDS PAGE and the protein concentration was determined by measuring the absorbance at 280 nm by using a Nanodrop spectrophotometer and using a calculated molar extinction coefficient of 102,680 M^-1^ cm^-1^. Purified protein was aliquoted and frozen at -80 °C until use.

### Purification of CHIKV active-site derivatives

All nsP2 active-site derivatives were purified initially using a rapid nickel affinity spin-column protocol (Qiagen) for biochemical screening experiments. For a subset of variants (S193A, K192L, and K192R), proteins were additionally subjected to full-scale purification, including SUMO cleavage and ion-exchange chromatography, using the same procedure as described for wild-type nsP2. Frozen cell pellets were thawed on ice and resuspended in lysis buffer (50 mM sodium phosphate pH 7.8, 500 mM NaCl, 1 mM TCEP, 20% glycerol) at 5 mL buffer per gram of cells. Cells were lysed by high-pressure homogenization as described for wild-type nsP2. NP-40 and PEI were added to final concentrations of 0.1% (v/v) and 0.25% (v/v), respectively, followed by stirring and clarification by centrifugation at 70,000 × g for 30 minutes at 4 °C. The clarified supernatant was subjected to ammonium sulfate precipitation at 40% saturation. Precipitated protein was collected by centrifugation at 70,000 × g for 30 minutes, resuspended in buffer A (25 mM HEPES pH 7.5, 500 mM NaCl, 20% glycerol, 1 mM TCEP with 10 mM imidazole), and loaded onto Ni-NTA spin columns (QIAGEN, #31014) equilibrated in buffer A. Columns were washed with buffer A containing 10 mM and 50 mM imidazole, and proteins were eluted with buffer A containing 500 mM imidazole. Eluted proteins were used directly for biochemical assays. Proteins purified by nickel spin columns are shown in Fig. 1. Select active-site derivatives (K192L, K192R, S193A) were subsequently purified with the full purification protocol described above for use in Fig. 2 experiments.

**Figure 1.**
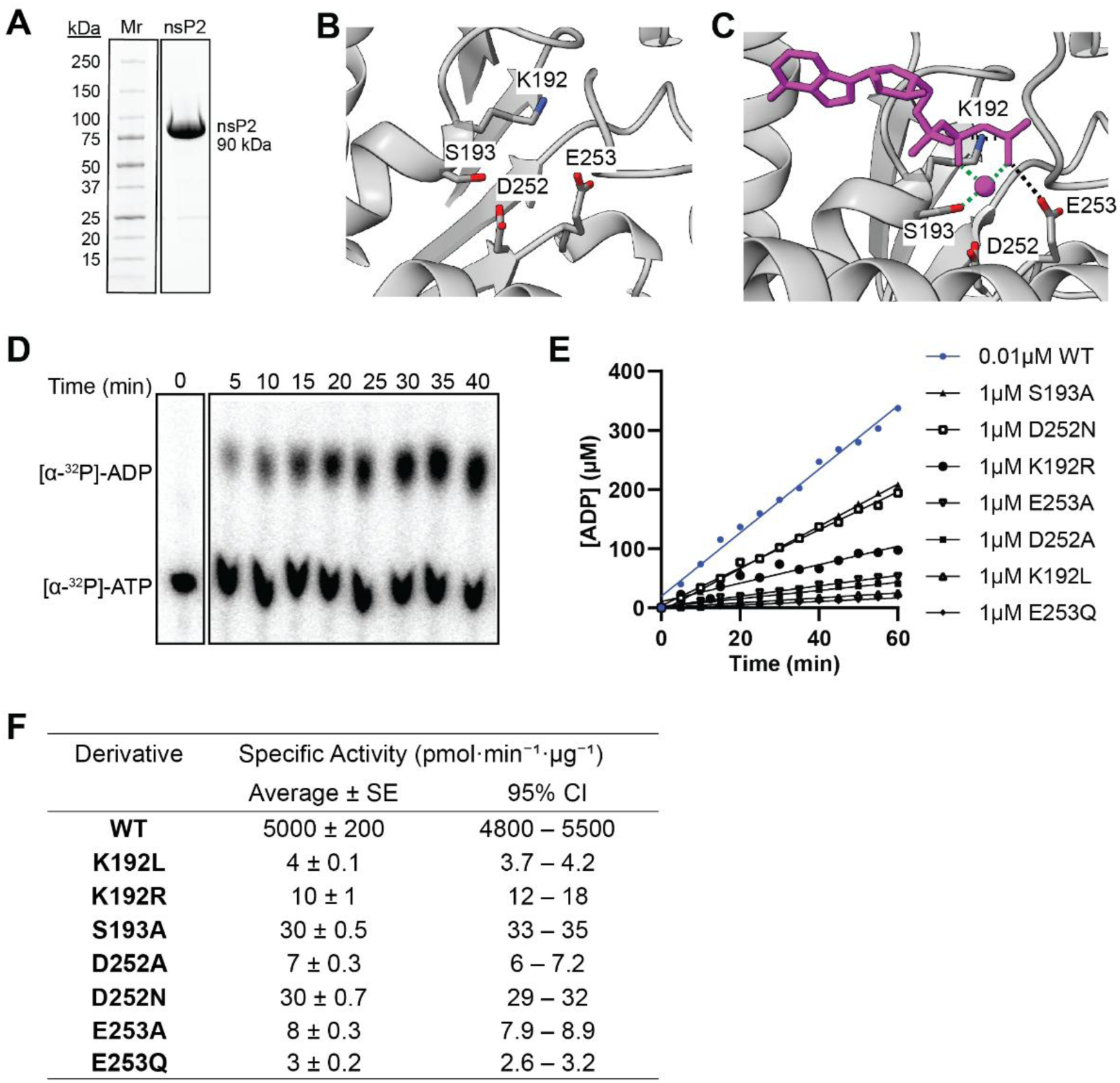
Purification and ATPase specific activity of CHIKV nsP2 ATPase active-site derivatives. **(A**) SDS-PAGE analysis of bacterially expressed, purified CHIKV nsP2. 13 µg of purified nsP2 at 95% purity is shown. Broad molecular weight markers (Mr) and corresponding molecular weights are indicated. Full expression and purification of recombinant wild type nsP2 are outlined in Fig. S1. **(B-C)** AlphaFold3 models of the nsP2 ATPase active site generated without (B) or with (C) bound ATP and Mg²⁺. Walker A (K192, S193) and Walker B (D252, E253) residues are highlighted. Hydrogen-bonding (black) and metal-coordinating (green) interactions are indicated. **(D)** ATP hydrolysis by wild-type nsP2 monitored by thin-layer chromatography. 0.01 µM nsP2 was mixed with 1 mM ATP and conversion of ATP to ADP was monitored. The TLC plate shown is one representative wild-type experiment from the quantification in (E/F). **(E)** ATPase activity of nsP2 wild-type and active-site derivatives purified using Ni-NTA spin columns. Column elutions containing SUMO-nsP2 proteins were used for ATPase reactions which contained 1 µM enzyme (0.01 µM for wild type) and 1 mM ATP. Reactions were quenched at various time points and resolved via TLC. Rates were determined by linear regression of ADP formation. **(F)** ATPase specific activities for nsP2 wild-type and derivatives. Rates from panel (E) (slope = µM ADP·min⁻¹) were converted to specific activity (pmol ADP·min⁻¹·µg⁻¹ nsP2) using the enzyme concentration and molecular weight of SUMO-nsP2. Values are reported as mean ± SE with 95% confidence intervals.

**Figure 2.**
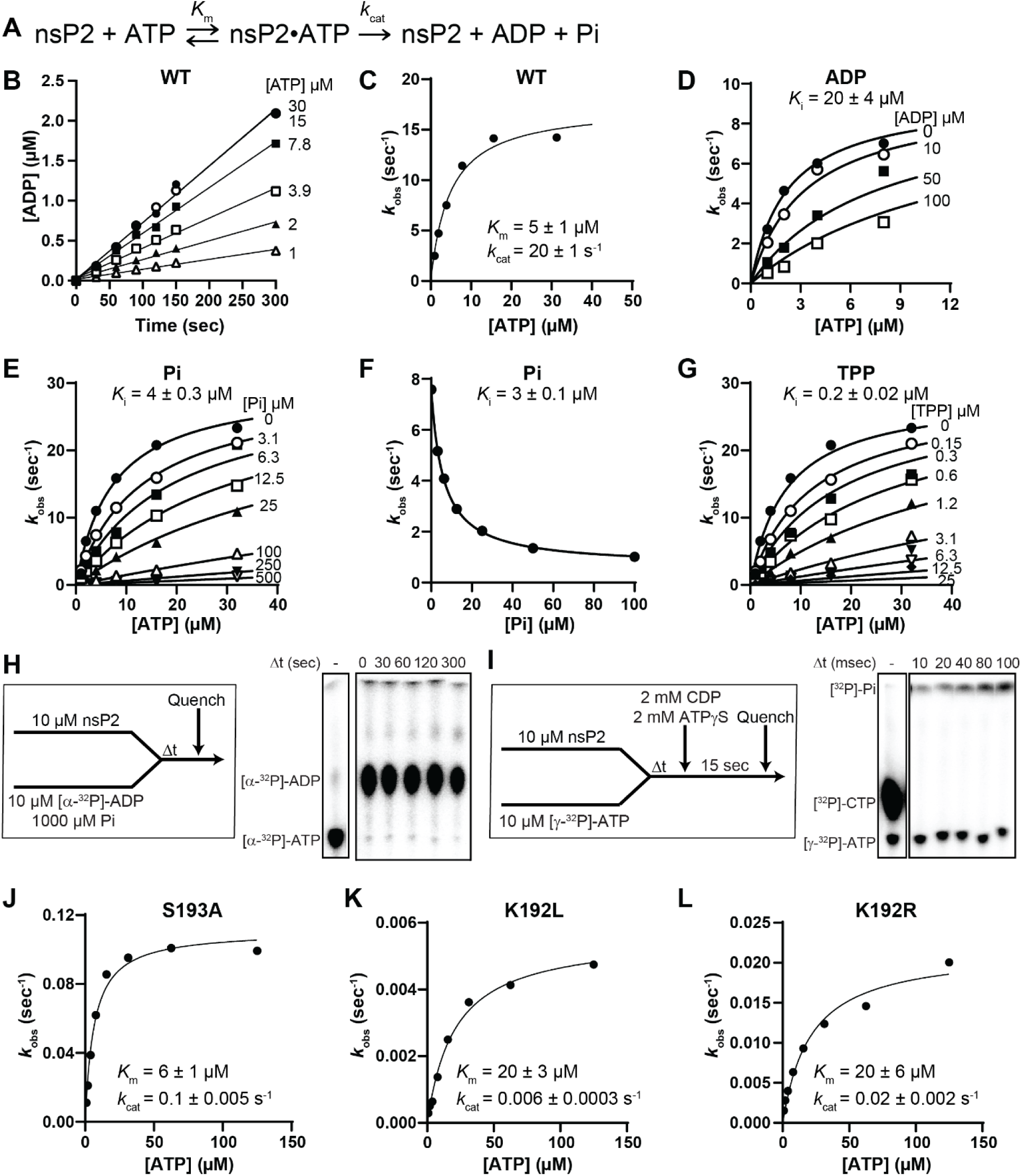
Steady-state ATP hydrolysis and product inhibition by CHIKV nsP2. **(A)** Mechanistic scheme for steady-state ATP hydrolysis by nsP2. **(B)** Initial rates of ATP hydrolysis were measured for WT nsP2 (0.5 nM) across a range of ATP concentrations under steady-state conditions. **(C)** Initial velocities from (B) were converted to observed rate constants (*k*_obs_, s^-1^) by normalizing to enzyme concentration (0.5 nM). The resulting *k*_obs_ values were plotted as a function of ATP concentration and fit to the Michaelis-Menten equation by nonlinear regression using all measured ATP concentrations to determine *K*_*m*_ and *k*_cat_ values. Data are shown as mean ± standard error. **(D-E)** Product inhibition of nsP2 steady-state ATPase activity by ADP (D) or inorganic phosphate (Pi) (E). Reactions were performed using varying ATP concentrations (0-32 µM) and fixed concentrations of ADP (0-100 µM) or Pi supplied as potassium phosphate (0-250 µM). Initial velocity data were globally fit by nonlinear regression. Competitive inhibition models best described the data, alternative inhibition models are shown in Fig. S2. **(F)** Effect of inorganic phosphate on nsP2 steady-state ATPase activity measured at a fixed ATP concentration equal to *K*_*m*_ (4 µM). Reactions were performed under the same conditions as in panels D-E, with Pi concentrations ranging from 0-100 µM. Data were fit to an inhibitor-versus-response model to determine an IC₅₀ value, which was converted to an inhibition constant (*K*_*i*_) via the Cheng-Prusoff equation. **(G)** Product inhibition of nsP2 steady-state ATPase activity by tripolyphosphate (TPP) measured under the same experimental conditions as in panels D-E. Initial velocity data collected in the presence of increasing concentrations of TPP (0-25 µM). Competitive inhibition provided the best-constrained fit and is shown here; alternative inhibition models are presented in Fig. S2. **(H-I)** Assessment of ATP synthesis by nsP2 in the reverse reaction under steady-state and pre-steady-state conditions. **(H)** Steady-state ATP synthesis assay where nsP2 (10 µM) was incubated with ADP (10 µM) and inorganic phosphate (1 mM). **(I)** Pre-steady-state ATP synthesis experiment in which nsP2 (10 µM) was rapidly mixed with ATP (10 µM), followed by addition of excess CDP (mM) and TPγS (mM). **(J-L)** Michaelis-Menten ATPase activity plots of nsP2 active-site derivatives K192L, K192R, and S193A measured using 0.5 µM enzyme. Initial velocities were obtained from linear regression from a 5 minutes time point for K192L and K192R and from a 2.5 minute time point for S193A (data not shown), converted to observed rate constants (*k*_obs_, s^-1^) by normalizing to the enzyme concentration (0.5 µM), and were fit by nonlinear regression to determine *K*_*m*_ and *k*_*cat*_ values. Reaction conditions were as in panel B.

### Steady-state NTPase assays

Reactions contained 25 mM HEPES pH 7.5, 1 mM magnesium acetate, 50 mM potassium glutamate, 1 mM TCEP, and 0.025 µCi/µL [α^32^P]-NTP/dNTP. Specific concentrations of substrate or enzyme, along with any deviations from the above, are indicated in the appropriate figure legend. Reactions were initiated by the addition of nsP2 and incubated at 30 °C. Reactions were quenched at various times by addition of EDTA to 250 mM final. The volume of enzyme added to any reaction was always less than or equal to one-tenth the total volume. Products were resolved by thin-layer chromatography (TLC). Initial velocities were calculated using linear regression and converted to observed rate constant, (*k*_obs_, s^-1^) by normalizing to nsP2 concentration. Data were fit by nonlinear regression to the Michaelis-Menten equation, where *k*_*cat*_ is the catalytic rate constant and *K*_*M*_is the Michaelis constant.

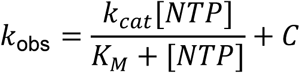

Specific activity was calculated from initial rates measured under steady-state conditions at saturating ATP concentrations. Initial velocities were determined from the linear portion of product formation time courses and converted to rates of ADP formation (pmol·min⁻¹) based on the total reaction volume and substrate specific activity. Rates were normalized to the amount of enzyme present in each reaction and reported as specific activity in units of pmol ADP·min⁻¹·µg⁻¹ nsP2.

### ATPase product inhibition assays

Steady-state ATPase product inhibition assays were performed to assess the effects of ADP, inorganic phosphate (Pi), and tripolyphosphate (TPP) on nsP2 ATP hydrolysis. Reactions contained 25 mM HEPES (pH 7.5), 1 mM magnesium acetate, 100 mM potassium glutamate, 0 mM TCEP, and 0.05 µCi/µL [α-³²P]-ATP. Reactions were conducted using 0.5 nM nsP2 and varying concentrations of ATP (0-32 µM) in the presence of fixed concentrations of ADP (0-100 µM), Pi supplied as potassium phosphate (pH 7.5, 0-250 µM), or TPP (0-25 µM), as indicated in the figure legends. Reactions were initiated by addition of nsP2 and incubated at 30 °C for 2 minutes before quenching with EDTA to a final concentration of 250 mM. The volume of enzyme added to each reaction did not exceed one-tenth of the total reaction volume. Reaction products were resolved by TLC. Initial velocities were determined by linear regression of ADP formation and converted to observed rate constants (*k*_obs_, s^-1^) by normalization to the nsP2 concentration. The resulting *k*_obs_ values were globally fit by nonlinear regression to competitive, noncompetitive, and uncompetitive inhibition models, as indicated below, using GraphPad Prism.

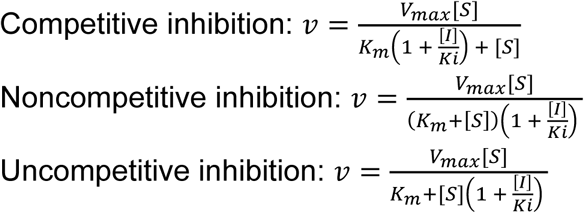

### Steady-state and pre-steady-state reverse ATPase reaction

ATP synthesis by nsP2 in the reverse reaction was assessed under both steady-state and pre-steady-state conditions. Steady-state ATP synthesis assays were performed using a chemical quench-flow apparatus (Kintek), in whi h nsP2 (10 µM) was incubated with [α-³²P]-ADP (10 µM) and inorganic phosphate (1 mM). Reactions were monitored for up to 300 s. Reaction products were resolved and visualized by TLC.

Pre-steady-state ATP synthesis experiments were conducted using a chemical quench-flow apparatus by rapidly mixing nsP2 (10 µM) with [γ-³²P]-ATP (10 µM), followed by the addition of excess CDP (2 mM) and ATPγS (2 mM) to prevent hydrolysis of any newly formed nucleotide. Reactions were monitored over a 100 ms time course, and reaction products were resolved and visualized by TLC.

### Pulse-quench and pulse-chase-quench experiments

Pulse-quench and pulse-chase-quench experiments were performed using a chemical quench-flow apparatus (Kintek). nsP2 (10 µM) was rapidly mixed with [α-³²P]-ATP (1 µM) and quenched with HCl at the indicated time points. For pulse-chase-quench experiments, which allow accumulation of internally trapped *nsP2·ATP prior to quenching, nsP2 (10 µM) was rapidly mixed with [α-³²P]-ATP (1 µM), followed at the indicated time points by rapid mixing with excess unlabeled ATP (1 mM). After an additional 30 s incubation, reactions were quenched with HCl. Reaction products were analyzed by TLC. Kinetic data were fit using kinetic modeling software (KinTek Explorer) to extract rate constants for the minimal mechanism.

### Malachite green phosphate binding assay

Inorganic phosphate (Pi) production was quantified using a malachite green colorimetric assay (Bioassay Systems), in which free Pi binds malachite green and produces an increase in absorbance at 620 nm. All assays were performed in 25 mM HEPES (pH 7.5), 1 mM magnesium acetate, 1 mM TCEP, and 100 mM potassium glutamate. A phosphate standard curve was generated to enable quantification of Pi produced in nsP2 reactions. Potassium phosphate (potassium phosphate, pH 7.5) standards ranging from 0 to 200 µM Pi were prepared under identical buffer conditions. Standard curve samples were mixed by combining 20 µL of each Pi standard with 80 µL malachite green reagent in non-binding 96-well plates. Samples were incubated for 30 minutes at room temperature, and absorbance at 620 nm was measured using a BioTek Synergy H1 plate reader. Absorbance values were fit by linear regression and used to convert experimental absorbance measurements to micromolar Pi concentrations.

To assess whether TPP is a substrate for nsP2, phosphate production was measured in reactions containing ATP or TPP. Reactions (100 µL total volume) contained ATP or TPP at the indicated concentrations (20, or 50 µM) and were initiated by addition of nsP2 to a final concentration of 20 nM. Reactions were incubated at 30 °C for either 5 or 30 minutes. Reactions were quenched by direct addition of malachite green reagent, with 20 µL of each reaction mixed immediately with 80 µL malachite green reagent. Samples were incubated for 30 minutes at room temperature prior to measuring absorbance at 620 nm. Phosphate concentrations were calculated using the phosphate standard curve.

### Assessment of ATPγS as a substrate for nsP2 using DP-Glo

ATPase activity was measured using the ADP-Glo™ assay (Promega), which quantifies ADP formation by enzymatic depletion of unreacted ATP followed by luciferase-based detection of ADP-derived ATP as a luminescent signal. Reactions (60 µL total volume) were performed in 25 mM HEPES (pH 7.5), 20 mM potassium glutamate, 0.5 mM magnesium acetate, and mM TCEP, using mM TP or TPγS and 50 nM nsP2. To terminate reactions, 10 µL of ADP-Glo™ Reagent was added to each sample, followed by incubation for 40 minutes at room temperature to deplete remaining ATP. Subsequently, 20 µL of Kinase Detection Reagent was added, and samples were incubated for an additional 1 h at room temperature to allow luminescent signal development. Following incubation, 40 µL of each reaction was transferred to a non-binding white 96-well plate, and luminescence was measured using a BioTek Synergy H1 plate reader. A nucleotide standard curve ranging from 0 to mM TP or TPγS was generated under identical buffer conditions to verify assay performance and signal responsiveness.

### Mant-ATP binding FRET assays

Equilibrium binding of mant-ATPγS to nsP2 was monitored using a tryptophan-to-mant FRET assay. Reactions were performed in 30 uL total volume in non-binding 384-well plates. To validate the tryptophan-to-mant FRET assay, fluorescence emission spectra were collected using a BioTek Synergy H1 plate reader equipped with a monochromator. Reactions contained nsP2 (1 µM) and mant-TPγS (0 µM) in binding buff r (5 mM HEPES, pH 7.5, mM magnesium acetate, 1mM TCEP, and 100 mM potassium glutamate). Spectra were collected for nsP2 alone, mant-ATPγS alone, and nsP2 + mant-ATPγS following excitation at 80 nm. Emission spectra were recorded from 300 to 500 nm. mant-ATPγS only spectra were used to define the background emission at 445 nm under 280-nm excitation.

Quantitative equilibrium binding measurements were performed using the same fluorescence setup. Reactions containing nsP2 (0.25 µM) were titrated with increasing concentrations of mant-ATPγS (0.00 -2.5 µM) in binding buffer. Samples were incubated for 10 minutes at room temperature to reach equilibrium. Fluorescence was measured using the BioTek Synergy H1 plate reader in monochromator mode, with excitation at 280 nm and emission monitored at 445 nm. For each mant-ATPγS concentration, mant-ATPγS only controls were collected under identical conditions and subtracted from the corresponding nsP2-containing samples to correct for background fluorescence. Baseline-subtracted fluorescence data were fit by nonlinear regression to a hyperbolic binding model to determine the apparent dissociation constant (*K*_d,app_). Specifically, data from mant-ATPγS titration experiments were fit to:

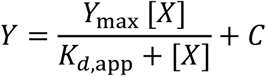

where [*X*] is the mant-ATPγS concentration, *Y* is the baseline-subtracted FRET fluorescence intensity at 445 nm, *Y*_max_ is the maximal signal at saturation and *K*_*d,app*_ is the apparent dissociation constant. Direct binding experiments were performed in triplicate (*n* = 3).

Competitive displacement of mant-ATPγS was performed to ass ss binding of unlabeled nucleotides and phosphate species. nsP2 (0.125 µM) was incubated with mant-ATPγS (0. µM) in binding buff r ontaining 5 mM HEPES (pH 7.5), 00 mM potassium glutamate, and 1 mM TCEP. Magnesium acetate was included at 1 mM for TPγS omp tition xp rim nts and at 5 mM for DP, inorgani phosphat (Pi), and tripolyphosphate (TPP) competition experiments. Increasing concentrations of competitors were added as indicated. Fluorescence was measured as described above, and data were normalized to percent relative fluorescence, with the signal in the absence of competitor defined as 100%. Data were fit by nonlinear regression to a four-parameter inhibition model. Apparent *IC*₅₀ values were converted to inhibition constants (*K*ᵢ) using a tight-binding form of the Cheng-Prusoff equation, where: [*L*]_50_= free labeled ligand concentration at the point where inhibition is 50%. It was calculated using a quadratic binding equilibrium assuming 1:1 binding between nsP2 and mant-ATPγS, using the independently measured *K*_*d*_.

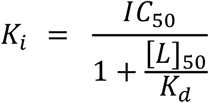

Reported *K*ᵢ values are best-fit estimates; standard errors were calculated from the 95% confidence intervals of the corresponding *IC*₅₀ fits. Exp rim nts with TPγS and DP were performed in triplicate (*n* = 3). Experiments with Pi and TPP were performed three times independently; independent experiments yielded similar results, one representative experiment is shown.

### Stopped-Flow kinetics of mant-nucleotide association and dissociation

Rapid kinetic measurements were performed using a Stopped-Flow SF-300x instrument (KinTek) equipped with a temperature-controlled water bath. All experiments were conducted at 30 °C. Binding and dissociation were monitored by tryptophan-to-mant fluorescence resonance energy transfer (FRET), which reports on formation or loss of nsP2·mant-TPγS complexes. Samples were excited at 280 nm, and fluorescence emission was monitored through a 460-nm long-pass filter (Chroma Technology, HQ440l) prior to photomultiplier detection. All reactions were performed in binding buffer containing 25 mM HEPES (pH 7.5), 1 mM TCEP, and potassium glutamate at 20 mM or 100 mM, as indicated. Magnesium acetate was included at a 1:2 molar ratio relative to nucleotide. Reactions were initiated by rapid 1:1 mixing. For each condition, at least three individual fluorescence traces were collected and averaged. All kinetic experiments were performed in triplicate (*n* = 3). Relative fluorescence intensity was plotted as a function of time and analyzed by nonlinear regression as described below.

Association and dissociation kinetics for mant-ATPγS were measured at potassium glutamate concentrations of 20 mM and 100 mM, in the absence or presence of the nsP2 inhibitor RA-NSP2-1 (15 µM), as indicated. mant-ATPγS association kinetics were measured by rapidly mixing mant-ATPγS (0.1 µM final) with nsP2 at concentrations ranging from 0.5-3 µM final under pseudo-first-order conditions.

Formation of the nsP2·mant-ATPγS complex was monitored as an in r as in sensitized mant fluorescence. Observed rate constants (*k*_obs_) were obtained by fitting fluorescence time courses to a single-exponential equation and replotted as a function of nsP2 concentration to determine second-order association rate constants (*k*_on_). Mant-ATPγS dissociation kinetics were measured by pre-forming nsP2·mant-ATPγS complexes (1 µM nsP2 and 10 µM mant-ATPγS final) and rapidly mixing with excess unlabeled TPγS (1 mM final) to prevent rebinding. Dissociation was monitored as a decrease in sensitized mant fluorescence and analyzed by nonlinear regression.

Association and dissociation kinetics for mant-ATP were measured only in the presence of RA-NSP2-1 (15 µM final) to suppress nucleotide hydrolysis and allow accurate measurement of binding kinetics. mant-ATP association and dissociation experiments were performed using the same experimental designs as described for mant-ATPγS, with mant-ATP substituted for mant-ATPγS. at onstants were determined at both 20 mM and 100 mM potassium glutamate.

### Classical molecular dynamics (cMD) simulations

Input structures were prepared by modeling the full-length nsP2 (1-798) with the command-line AlphaFold3 (18). Three systems were modeled: nsP2, nsP2:ATP-Mg^2+^, and nsP2:ATP-Mg^2+^:RA-NSP2-1. The MD simulation files were made by CHARMM-GUI using the AlphaFold3 models as input(19). The complexes were placed in the center of a cubic box such that the faces of the box are 1 nm apart from the protein. Boxes were solvated with TIP3P water and ionized with 150 mM KCl to afford an electroneutral system. Protein was parameterized by AmberFF14SB, and the ligands were parameterized by CGenFF (20).

The cMD simulations were carried out by GROMACS 2026.0 (21). Periodic boundary conditions were applied in all directions. Complexes were energy minimized by the Steepest Descent algorithm. The simulation box equilibrated to 298 K by running the system for 100 ps under the NVT ensemble while restraining the protein complex under harmonic potential with 400 kJ mol^-1^.nm^-2^ force constantly applied on the heavy atoms. The system was further equilibrated for 100 ps under the NPT ensemble with identical position restraints as the NVT equilibration. The production runs were carried out with 2 fs time step for a total 1 μs, and the frames were recorded every 100 ps. The temperature was controlled by the V-scale algorithm, and the pressure was controlled by the C-rescale algorithm. Covalent bonds involving hydrogen atoms were constrained with the P-LINCS method. Electrostatic interactions were treated by the smooth particle mesh Ewald method.

To monitor the dynamics of the protein backbone, root mean square fluctuation (RMSF) of the Cα atoms were calculated using the MDAnalysis package (22). Only the helicase domain of nsP2, residue 1 to 465, was considered for RMSF calculations.

### Accelerated molecular dynamics (aMD) simulations

Input structures of full length nsP2 and complexes with ATP-Mg2+ and inhibitor were prepared similar to those described for the cMD simulations. Each model was subjected to all-atom molecular dynamics simulations using AMBER software suite(23) with parameters from the amber14SB forcefield (24).

All-atom accelerated molecular dynamics simulations were performed in explicit water (TIP3P model) (25); a distance of 16 Å between the edge of the truncated octahedron solvent box and any protein atoms was imposed. A cutoff radius of 12 Å was used in the calculations of non-bonded interactions with periodic boundary conditions applied; particle mesh Ewald method (26,27) was used to treat electrostatic interactions. The systems were relaxed by two cycles of energy minimization and slow heating to 300 K using Particle Mesh Ewald Molecular Dynamics (PMEMD) under constant volume and temperature (NVT) conditions. Afterward, two cycles of constant pressure (NPT) dynamics were performed; the second cycle reducing the non-bonded cutoff to 9 Å both employing a 1 fs integration time-step for a total of 5 ns of simulation time. The equilibrated systems were then subjected to 10 ns of classical MD under NVT conditions with 1 fs integration time step. Finally, aMD was performed using the dual-boost strategy (28) for 140 ns with snapshots saved at 1 ps interval and 2 fs integration time step. Analyses of the trajectories from MD simulations were merged using CPPTRAJ (29) and analyzed according to the methods outlined below.

### Principal component analysis of collective motions

To characterize the dominant, correlated collective motions of the enzyme, Principal Component Analysis (PCA) was performed utilizing a combined MDTraj (v1.9) (30) and ProDy (v2.6.1) (31) pipeline. Trajectories were first stripped to backbone heavy atoms (N, CA, C, O) to preserve secondary structure geometry during visualization. The structural ensembles were then translationally and rotationally superposed onto the reference coordinates to eliminate global tumbling artifacts. Positional covariance matrices were constructed from the aligned atomic coordinates and diagonalized to extract the principal components (eigenvectors) and their corresponding variances (eigenvalues).

To effectively decouple localized allosteric mechanisms from large-scale, low-frequency domain breathing, PCA was computed at two distinct structural scales: (1) globally, across the entire helicase domain, and (2) locally, with the covariance matrix restricted exclusively to the backbone atoms of the interacting subdomains. The dominant eigenvectors representing the primary axes of motion (PC1) were exported and rendered as 3D vector projections (porcupine plots) mapped onto the protein backbone using the Normal Mode Wizard (NMWiz) plugin within Visual Molecular Dynamics (VMD) (32). Animations were created using the VMD Movie Maker plugin for each of the respective trajectories.

### 5’-32P-labeling of RNA substrates

RNA oligonucleotides were end-labeled by using [*γ*-^32^P] ATP and T4 polynucleotide kinase. Reaction mixtures, with a typical volume of 50 μL, contained 0.5 μM [*γ*-^32^P] ATP, 1 µM RNA oligonucleotide, 1× kinase buffer, and 0.4 unit/μL T4 polynucleotide kinase. Reaction mixtures were incubated at 37 °C for 60 minutes and then held at 65 °C for 5 minutes to heat inactivate T4 PNK.

### Annealing of dsRNA substrates

dsRNA substrates were produced by annealing 1 μM RNA oligonucleotides in 25 mM HEPES pH 7.5 and 50 mM NaCl in a Progene Thermocycler (Techne). Annealing reaction mixtures were heated to 90 °C for 1 minute and slowly cooled (5 °C/min) to 10 °C.

### RNA binding fluorescence polarization experiments

Increasing concentrations of nsP2 were mixed with 1 nM of duplexed fluorescein-labeled RNA in a binding buffer containing 25 mM HEPES, 1 mM magnesium acetate, 1 mM TCEP, 50 mM or 100 mM potassium glutamate at pH 7.5. Reactions were incubated at room temperature for 10 minutes in a black 384-well non-binding plate and then read with a dual wavelength Ex: 485/20 Em: 528/20 fluorescence polarization cube using a Biotek Synergy H1 plate reader. Data from protein titration experiments were fit to a hyperbola:

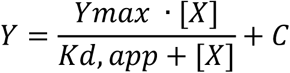

where X is the concentration of protein, Y is degree of polarization, *K*_d,app_ is the apparent dissociation constant, and Y_max_ is the maximum value of Y.

For stoichiometric binding experiments, nsP2 was titrated into a fixed concentration of RNA present in >10x molar excess to the determined *K*_d,app_ (50 or 150 nM, for U40 and U20 RNA substrates, respectively). Fluorescence polarization values were plotted as a function of total nsP2 concentration and fit by linear regression in the pre- and post-saturation regions. The intersection of these two linear fits corresponds to the molar ratio of nsP2 to RNA at saturation and was used to determine binding stoichiometry.

### RNA helicase assays

Forked duplex RNA substrates were prepared by annealing a radiolabeled loading strand to an unlabeled displaced strand to generate partially duplex RNAs containing a single-stranded 5′ loading r gion, a ntral duplex region, and a 3′ single-stranded tail. Substrates contained either a U20 or U40 5′ single-stranded region, a 9- or 7-bp duplex region, and a 3′ U10 tail. RNA sequences are listed in Fig. 9A. An unlabeled RNA trap strand was included in unwinding reactions to prevent re-annealing of unwound products. Multiple-turnover RNA unwinding reactions contained 25 mM HEPES (pH 7.5), 20 mM potassium glutamate, 1 mM TCEP, 10 nM duplex RNA substrate, 0.5 µM RNA trap strand, 2 mM ATP, and 1 mM magnesium acetate, unless otherwise indicated. Reactions were initiated by addition of nsP2 to a final concentration of 250 nM and incubated at 30 °C. At the indicated time points, reactions were quenched by 1:1 mixing with 2× quench buffer, yielding final concentrations of 100 mM EDTA, 0.33% (w/v) SDS, 1 mM poly(rU) (UMP equivalents), 0.025% (w/v) bromophenol blue, 0.025% (w/v) xylene cyanol, and 5% (v/v) glycerol. Quenched samples were analyzed by native PAGE. Gels were visualized by phosphorimaging, and ssRNA product formation was quantified. Data are reported as ssRNA concentration versus time. Kinetic fitting was not performed under these conditions due to ATP depletion during the course of the reaction (see Fig. S4).

Optimization experiments examining magnesium dependence, ATP dependence, and nucleotide specificity were performed under otherwise identical conditions, except that 10 mM TCEP was used. Magnesium acetate and ATP concentrations were varied as indicated in Fig. S3. For experiments maintaining a constant Mg²⁺:ATP ratio, ATP and magnesium acetate were varied together at a 1:2 molar ratio. Nucleotide specificity experiments were performed using ATP, CTP, GTP, or UTP under optimized conditions.

Single-turnover RNA unwinding assays were performed under conditions matching those used for multiple-turnover reactions. Reactions contained 25 mM HEPES (pH 7.5), 20 mM potassium glutamate, 1 mM TCEP, 10 nM duplex RNA substrate, and 250 nM nsP2. nsP2 was pre-incubated with RNA for 5 minutes at 30 °C prior to initiation of unwinding. Unwinding was initiated by rapid addition of 2 mM ATP, 1 mM magnesium acetate, 0.5 µM RNA trap strand, and 100 µM heparin. Heparin was included to prevent rebinding of nsP2 following strand separation. Reactions were incubated at 30 °C and quenched at the indicated time points by 1:1 mixing with 2× quench buffer as described above. Products were resolved by native PAGE and visualized by phosphorimaging. Single-turnover unwinding kinetics were quantified by measuring ssRNA product formation over time. Data were fit by nonlinear regression to a single-exponential model to obtain apparent unwinding rate constants (*k*_obs_) using the following equation: *Y(t) = A · (1 − e^−kobs·t^)* Reported values represent mean ± SD from two independent experiments (*n* = 2).

### ATP regeneration systems

ATPase activity assays were performed with nsP2 (20 nM) in 25 mM HEPES (pH 7.5), 1 mM TCEP, 1 mM magnesium acetate, and 1 mM ATP at 30 °C. ATP regeneration systems were included as indicated: (i) creatine kinase (0.05 mg/mL) with creatine phosphate (20 mM), (ii) pyruvate kinase (0.05 mg/mL) with phosphoenolpyruvate (20 mM), or (iii) a commercial ATP regeneration system (Enzo, 1×). Reactions were initiated by addition of nsP2. Aliquots were removed at 5, 10, and 15 minutes, quenched by addition of an equal volume of 500 mM EDTA, and 1 µL of each sample was spotted onto PEI-cellulose TLC plates. Products were resolved using 0.3 M potassium phosphate buffer (pH 7.0) and visualized by autoradiography.

### RNA unwinding assays with ATP regeneration systems

Multiple-turnover RNA unwinding assays were performed using the U20-ds9-U10/ds9-U10 substrate (10 nM) in 25 mM HEPES (pH 7.5), 1 mM TCEP, 5 mM magnesium acetate, 5 mM ATP, 20 mM potassium glutamate, 0.5 µM RNA trap strand, and nsP2 (50 nM). Reactions were incubated at 30 °C in the presence of either creatine kinase with creatine phosphate (5 or 20 mM) or the commercial ATP regeneration system (Enzo). Reactions were quenched with the addition of 250 mM EDTA final. Reaction products were resolved on native polyacrylamide gels and visualized as described above.

### Product analysis: TLC

1 µl of the quenched reaction was spotted onto polyethyleneimine-cellulose TLC plates (EM Science). TLC plates were developed in 0.3 M potassium phosphate, pH 7.0, dried and exposed to a Phosphorimager screen. TLC plates were visualized by using a Phosphorimager and quantified by using the ImageQuant software (Cytiva) to determine the amount of NTP hydrolyzed to NDP. The amount of NDP was plotted as a function of time.

### Product analysis: native PAGE

10 µl of the quenched reaction mixtures was loaded on a 20% acrylamide, 0.53% bisacrylamide native polyacrylamide gel containing 1× TBE. Electrophoresis was performed in 1× TBE at 15 mA. Gels were visualized by using a phosphorimager and quantitated by using the ImageQuant software (Molecular Dynamics) to determine the amount of strand separation. The fraction of RNA that had been unwound by the helicase was corrected for the efficiency of the trapping strand in preventing reannealing of products. This correction factor was determined for each experiment by heating a sample of the RNA substrate at 95 °C for 1 minute then cooled to 10 °C at a rate of 5 °C/min in the presence of the trapping strand. The quantity of unwinding substrate that was prevented from reannealing was typically ∼98%. The amount of RNA unwound was plotted as a function of time.

### Data analysis

All gels shown are representative, single experiments that have been performed at least three to four individual times to define the concentration or time range shown with similar results. In all cases, values for parameters measured during individual trials were within the limits of the error reported for the final experiments. Data were fit by either linear or nonlinear regression using the program GraphPad Prism v7.03 (GraphPad Software Inc.)

## Results

### Expression and Purification of Wild-type Chikungunya Virus Non-structural Protein 2 and Derivatives with Substitutions in Conserved Active-site Residues

We produced proteins employed in this study in Escherichia coli using a vector expressing non-structural protein 2 (nsP2) as a fusion with an affinity-tagged small ubiquitin-like modifier (SUMO) protein (**Fig. S1A**). nsP2 accumulated to high levels in E. coli (**Fig. S1B**), with a substantial portion of the protein present in the soluble fraction of the cell lysate (**Fig. S1C**). We used a combination of Ni^2+^ affinity chromatography (**Fig. S1D**) and ion exchange chromatography (**Fig. S1E**) to isolate nsP2 (**Fig. 1A**). We removed the affinity tag by cleavage with the SUMO carboxy-terminal hydrolase, Ulp1, after elution from the Ni^2+^ column (compare lane Ni Pool to lane Ni Pool + Ulp1 in **Fig. S1E**). The purification protocol is quite robust and reproducible (**Fig. S1F**). Derivatives used in Fig. 1 were purified using a Ni²⁺ spin column, whereas selected variants (K192L, K192R, S193A) analyzed in Fig. 2 were purified under the same conditions as wild-type protein.

### Interrogation of the Conserved Residues in the Walker A and B Motifs

All superfamily 1 helicases use the “P-loop fold” comprised of two conserved motifs: Walker A and Walker B (33,34). nsP2 is no different. The Walker A motif of nsP2 includes Lys-192 and Ser-193, and the Walker B motif includes Asp-252 and Glu-253 (**Fig. 1B**). These residues bind and organize the tripolyphosphate (TPP) moiety of the substrate nucleotide for hydrolysis (**Fig. 1C**). The snapshot shown in **Fig. 1C** derives from AlphaFold3 as all structures of SF1B helicases illustrating a catalytically competent state with nucleotide employ non-hydrolysable analogs (14,35,36). The chemical mechanism for P-loop enzymes has yet to be elucidated. Recently, Kozlova et al. suggested that the Walker-A serine and the Walker-B aspartate form a low-barrier hydrogen bond that promotes formation of a serine alkoxide that activates the catalytic water molecule by abstracting a proton (33). If this hypothesis is correct, then changing Ser-193 in our system to alanine (S193A) should inactivate the ATPase. Such a derivative could prove useful for binding and structural studies employing a nucleotide substrate instead of a non-hydrolysable analog.

We assayed TPas a tivity using an [α-^32^P]-labeled nucleoside triphosphate substrate and monitor formation of the [α-^32^P]-labeled nucleoside diphosphate product by thin-layer chromatography (**Fig. 1D**). A variety of solution conditions were evaluated and sensitivity of the reaction to chloride ions was noted (data not shown). Optimal conditions were: 25 mM HEPES (pH 7.5), 1 mM magnesium acetate, 20-100 mM potassium glutamate, and 1 mM TCEP. Product formation was linear over a 60-minute period of time (**Fig. 1E**), yielding a specific activity for WT nsP2 of 5000 ± 200 pmol·min^-1^·µg^-1^ (**Fig. 1F**). The activity of each derivative was reduced 100- to 1000-fold relative to WT (**Fig. 1E,F**); however, each retained sufficient residual ATPase activity to prevent their use in biochemical and/or biophysical experiments with natural substrates.

### Steady-state Kinetic Analysis

#### Steady-state parameters

The anticipated scheme for nsP2-catalyzed hydrolysis of ATP is shown in **Fig. 2A**. We assume the reaction to be irreversible as is typical for this class of enzymes. We monitored the kinetics of ADP formation over a range of concentrations of ATP (**Fig. 2B**). A plot of the observed rate (*k*_obs_) of ATP hydrolysis as a function of ATP concentration yielded values for the *K*_m_ and *k*_cat_ of 5 ± 1 µM and 20 ± 1.0 s^-1^, respectively (**Fig. 2C** and **Table 1**).

**Table 1.** Effects of Walker A motif substitutions on steady-state kinetics of nsP2. Steady-state ATPase kinetics were determined for wild-type nsP2 and Walker A motif derivatives using 0.5 nM wild-type enzyme or 0.5 µM derivative proteins and varying ATP concentrations. Reactions were performed at 30 °C in the presence of 1 mM magnesium acetate and 50 mM potassium glutamate, and ATP hydrolysis was monitored by TLC using [α-³²P]-ATP. Observed rate constants were fit to the Michaelis-Menten equation to determine *K*_*m*_ (µM) and *k*_*cat*_ (s⁻¹), reported as mean ± standard error with associated 95% confidence intervals. Catalytic efficiencies (*k*_*cat*_/*K*_*m*_) were calculated and reported in µM⁻¹·s⁻¹.

| Derivative | $K_m$ ($\mu\text{M}$) | | $k_{cat}$ ($\text{s}^{-1}$) | | $k_{cat}/K_m$ ($\mu\text{M}^{-1}\cdot\text{s}^{-1}$) | |
| --- | --- | --- | --- | --- | --- | --- |
| | Average ± SE | 95% CI | Average ± SE | 95% CI | Average ± SE | 95% CI |
| **WT** | 5 ± 1 | 3 - 8 | 17 ± 1.0 | 15 - 20 | 4 ± 0.9 | 2 - 6 |
| **K192L** | 20 ± 3.0 | 16 - 28 | 0.0112 ± 0.0005 | 0.010 - 0.012 | 0.00054 ± 0.00008 | 0.0004 - 0.0007 |
| **K192R** | 20 ± 6.3 | 12 - 37 | 0.022 ± 0.002 | 0.02 - 0.03 | 0.0010 ± 0.0003 | 0.0004 - 0.0017 |
| **S193A** | 6 ± 1 | 5 - 9 | 0.22 ± 0.01 | 0.20 - 0.24 | 0.034 ± 0.006 | 0.02 - 0.05 |

#### Product release

To determine if a preferred order of product release exists, we monitored the kinetics of ATP hydrolysis in the presence of varying concentrations of ADP (**Fig. 2D**) and Pi (**Fig. 2E**). Both datasets fit better to a competitive model of inhibition (**Figs. 2D,E**) than a noncompetitive (**Figs. S2A,C**) or uncompetitive **(Figs. S2B,D)** model. The inhibition constant (*K*_i_) value for ADP was 20 ± 4.0 µM (**Fig. 2D**) and that for Pi was 4.0 ± 0.3 µM (**Fig. 2E**). These data are consistent with both ADP and Pi binding to free enzyme individually. Assuming the *K*_i_ is equivalent to *K*_d_ in this case, Pi would be released at a rate slower than ADP if the rate constants for association are equivalent. Because these estimated dissociation rates exceed the observed steady-state turnover rate (*k*_cat_ ≈ 0 s⁻¹), product release, particularly Pi release, is unlikely to be the sole rate-limiting step in steady-state ATP hydrolysis by nsP2.

#### Pi inhibition

While some P-loop motor proteins are inhibited by inorganic phosphate, this behavior is typically attributed to Pi binding to an ADP-bound enzyme state (4,37). To our knowledge, Pi has not been established as a physiologically relevant inhibitor of P-loop helicases. We therefore re-examined the sensitivity of nsP2 to Pi by titrating Pi at a fixed concentration of ATP of 4 µM (**Fig. 2F**). Under these conditions, we obtained a *K*_i_ value of 3.0 ± 0.1 µM. Because the intracellular concentration of Pi is generally in the millimolar range, this result suggests that the solute environment within the spherule in which nsP2 functions may differ substantially from that of the surrounding cytoplasm.

#### TPP inhibition

The ability of Pi to compete so effectively with binding of ATP suggested that a large proportion of the energetics of nucleotide binding is driven by the gamma phosphate and perhaps the beta and alpha phosphates as well. To test this possibility, we evaluated the sensitivity of nsP2-catalyzed hydrolysis to tripolyphosphate (TPP). First, we demonstrated that TPP is not hydrolyzed by nsP2 using a colorimetric assay for detection of Pi (**Fig. S2E**). Next, we evaluated TPP as an inhibitor of ATP hydrolysis, which revealed TPP as a competitive inhibitor with respect to ATP with a *K*_i_ value 0.2 ± 0.02 µM, more than an order of magnitude greater affinity than Pi (**Fig. 2G**). Fits to noncompetitive and uncompetitive inhibition models are shown in **Fig. S2F,G**. These data indicate that the triphosphate moiety makes a major contribution to the unusually tight nucleotide binding observed for nsP2.

#### Reverse reaction

We evaluated the reverse reaction using two different experimental designs. First, we monitorwd conversion of [α-^32^P]-ADP to [α-^32^P]-ATP in the presence of Pi (**Fig. 2H**). Formation of [α-^32^P]-ATP was not observed over a 300 s time course (**Fig. 2H**). In the second design, we attempted to trap labeled ^32^Pi produced by hydrolysis of [γ-^32^P]-ATP and then have that labeled transferred to CDP to form [γ-^32^P]-CTP. ATPγS was present in the reaction to prevent any [γ-^32^P]-CTP formed during the experiment from being hydrolyzed (**Fig. 2I**). Over the 100 ms time course, none of the ^32^Pi formed was converted to [γ-^32^P]-CTP (**Fig. 2I**). We conclude that ATP hydrolysis is irreversible under the experimental conditions used here.

#### Walker-A motif and catalysis

Residues of the Walker-A motif are thought to influence ATP binding and perhaps stabilize the catalytically competent orientation of the TPP (33). Therefore, we anticipated that perturbation of this motif would cause a greater effect on *K*_m_ than on *k*_cat_. Moreover, changing Lys-192 to Leu should be more destabilizing to the catalytically competent state than an Arg at this position. In all cases, *k*_cat_ was impacted to the greatest extent (**Fig. 2J-L** and **Table 1**). Even more unexpected was the observation that the K192L and K192R derivatives exhibited nearly identical steady-state parameters (**Fig. 2K,L** and **Table 1**).

#### Nucleotide substrate specificity

We did not perform a complete steady-state kinetic analysis for all possible nucleotide substrates. We evaluated each at the *K*_m_ value for ATP (4 µM) and at saturating concentrations (1 mM). The specific activities measured were identical for all nucleotides evaluated, consistent with the terminal Pi or TPP driving binding (**Table 2**).

**Table 2.** Nucleotide substrate specificity of nsP2 under steady-state ATPase conditions. Steady-state ATPase and dNTPase activities of wild-type nsP2 were measured using 0.5 nM nsP2 and 4 μM nucleotide or 20 nM nsP2 and 1 mM nucleotide, as indicated. Activities are reported as specific activity (pmol ADP·min⁻¹·μg⁻¹ nsP2) calculated from initial rates. All reactions were performed in the presence of 1 mM magnesium acetate and 50 mM potassium glutamate. Values are reported as mean ± standard error with associated 95% confidence intervals.

| Nucleotide | Specific Activity ($\text{pmol} \cdot \text{min}^{-1} \cdot \mu\text{g}^{-1}$) | | | |
| --- | --- | --- | --- | --- |
| | 0.5 nM nsP2, 4 μM NTP | | 20 nM nsP2, 1 mM NTP | |
| | Average ± SE | 95% CI | Average ± SE | 95% CI |
| **ATP** | 900 ± 80 | 800 - 1000 | 6000 ± 700 | 4000 - 7000 |
| **CTP** | 600 ± 70 | 500 - 700 | 5000 ± 400 | 5000 - 6000 |
| **GTP** | 700 ± 40 | 600 - 800 | 7000 ± 900 | 5000 - 8000 |
| **UTP** | 700 ± 30 | 700 - 800 | 5000 ± 900 | 4000 - 7000 |
| **dATP** | 900 ± 70 | 800 - 1000 | 6000 ± 1000 | 4000 - 8000 |
| **dCTP** | 900 ± 100 | 700 - 1000 | 5000 ± 1000 | 3000 - 7000 |
| **dGTP** | 900 ± 90 | 700 - 1000 | 5000 ± 600 | 4000 - 6000 |
| **dTTP** | ND | ND | 7000 ± 2000 | 3000 - 10000 |

### Equilibrium Binding of Nucleotides

The steady-state kinetic analysis suggests that nsP2 binds to ATP much more tightly than the typical SF1 family member. Of course, *K*_m_ is not a dissociation constant. Therefore, we valuated nucleotidebinding directly using ’(3’)-O-(N-Methyl-anthraniloyl)-conjugated ATPγS (mant-ATPγS) as don in thepast for other enzymes (**Fig. 3A**) (37-39). B for moving forward, we demonstrated that ATPγS was not a substrate for nsP2, unlike some ATPases (**Fig. 3B**) (40).

**Figure 3.**
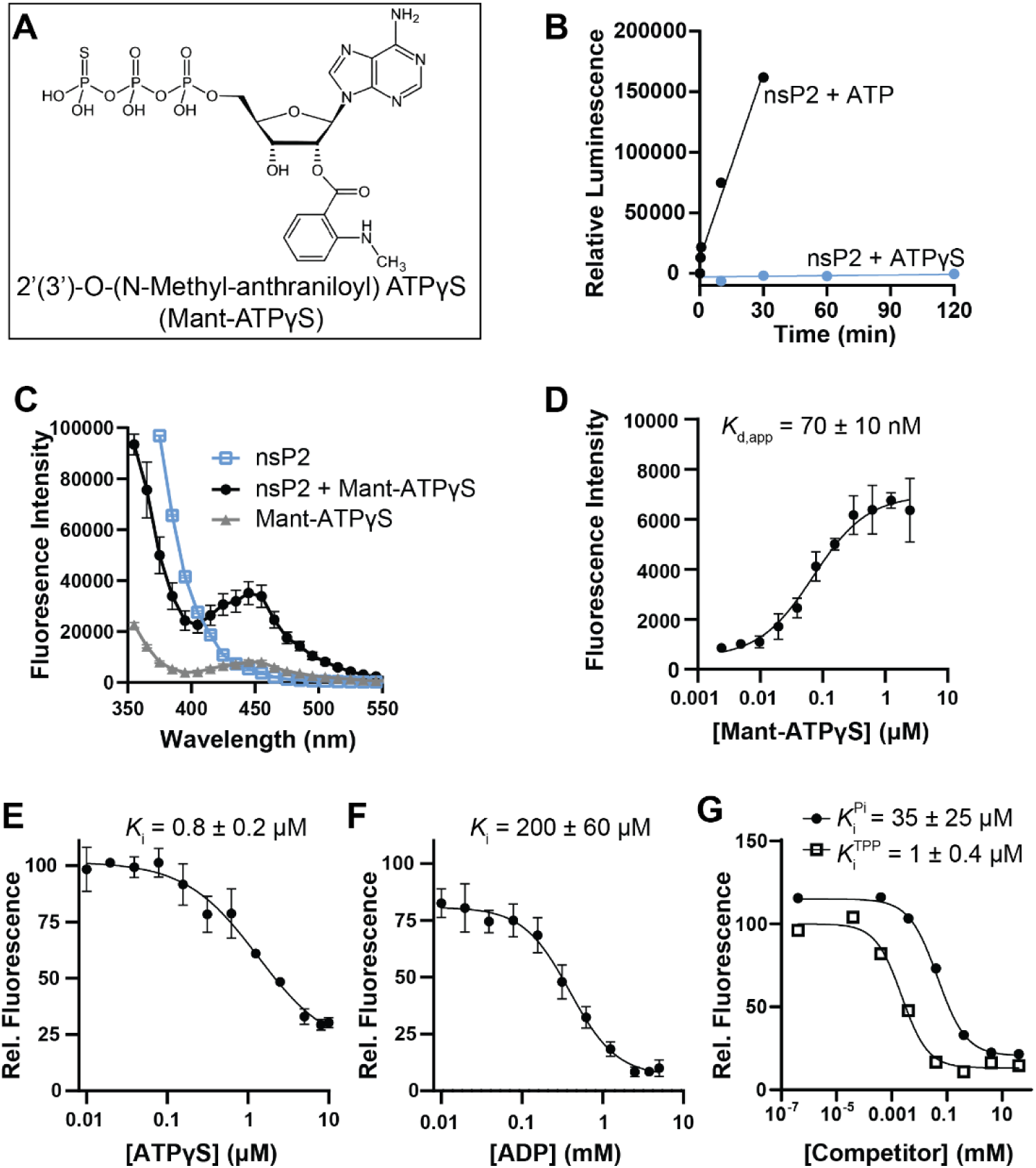
Equilibrium and competitive binding of Mant-ATPγS to nsPS2. **(A)** Chemical structure of ′(3′)-O-(N-methyl-anthraniloyl) ATPγS (mant-ATPγS). **(B)** ATPγS is not hydrolyzed by nsP2. Under the conditions tested, 50 nM nsP2 converted ∼80% of 1 mM ATP to ADP within 30 minutes, whereas no detectable hydrolysis of 1 mM ATPγS was observed after 120 minutes. No luminescence signal was detected in reactions containing ATP or ATPγS in the absence of enzyme (data not shown). **(C)** Representative tryptophan to mant FRET emission spectra collected using excitation at 80 nm. nsP2 alone (1 μM) exhibits an emission peak at 350 nm, whereas mant-ATPγS alone (10 μM) shows weak emission at 445 nm under 280-nm excitation. Addition of mant-ATPγS to nsP2 products an increase in 445-nm emission, consistent with FRET arising from formation of the nsP2·mant-ATPγS complex. Data in panels D-G were generated by subtracting mant-ATPγS-only emission at 445 nm from spectra collected in the presence of nsP2. **(D)** Direct binding of mant-ATPγS to nsP2. ns2P (0. 25 μM) was titrated with 0.002 5 μM mant-ATPγS. Data represent mean ± SD (*n* = 3). **(E-G)** Competitive binding experiments. nsP2 (0. 5 μM) was incubated with 0.1 μM mant-ATPγS and increasing concentrations of unlabeled competitor. ATPγS (E; 0-10 μM), ADP (F; 0-9 mM), or inorganic phosphate (Pi) and tripolyphosphate (TPP) (G; 0-40 mM) were added as indicated. Fluorescence data in panels E and F were normalized to percent relative fluorescence, with the signal in the absence of competitor defined as 100%. Data were fit by nonlinear regression, and IC₅₀ values were converted to inhibition constants (*K*_i_) using the Cheng-Prusoff equation.

Mant-ATPγS is fluorescent, with an excitation wavelength of 350 nm and an mission wavelength of 450 nm (38). Because tryptophan fluorescence of proteins emits at 350 nm when excited at 280 nm, fluorescence resonance energy transfer (FRET) can be used to illuminate only the bound mant-ATPγS (38). FRET from nsP2 to ATPγS was observed and produced a robust signal (**Fig. 3C**).

Titration of nsP2 with mant-ATPγS yielded a binding isotherm consistent with a *K*_d,app_ value of 70 ± 10 nM (**Fig. 3D**). This value represents very tight binding. To determine if the mant substituent contributes to the observed binding affinity, we performed a comp tition experiment with unlabeled ATPγS. This experiment yielded a *K*_i_ (*K*_d_) of 0.8 ± 0.2 µM (**Fig. 3E**). The gamma phosphate of the nucleotide contributes substantially to nucleotide binding as the *K*_i_ (*K*_d_) for ADP was 200 ± 60 µM (**Fig. 3F**). Both Pi and TPP competed for binding of mant-ATPγS, yielding *K*_i_ (*K*_d_) values of 35 ± 25 µM and 1.0 ± 0.4 µM, respectively (**Fig. 3G**). These data are consistent with the TPP substituent of the nucleoside triphosphate being sufficient to explain the observed affinity of nsP2 for nucleotides.

### Kinetics of Nucleotide Binding

We were also able to use mant-ATPγS to monitor the kinetics of binding using a stopped-flow device. We rapidly mixed nsP2 at varying concentrations with mant-ATPγS, and the signal for mission at 445 nm was monitored after excitation at 80 nm (**Fig. 4A**). The experiment was performed under pseudo-first-order conditions in the presence of 100 mM potassium glutamate. Each nsP2 concentration evaluated yielded a time course for association (**Fig. 4B**). We then determined the observed rate constant for binding at each nsP2 concentration by fitting each time-course to a single-exponential equation. Finally, we plotted the observed rate as a function of nsP2 concentration (**Fig. 4C**). The rate constant for association (*k*_on_) was determined from the slope of the line, which yielded a value of 115 ± 5 µM^-1^ s^-1^. Reducing the concentration of potassium glutamate to 20 mM caused an increase in the rate constant for association to 325 ± 30 µM^-1^ s^-1^ (**Table 3**).

**Figure 4.**
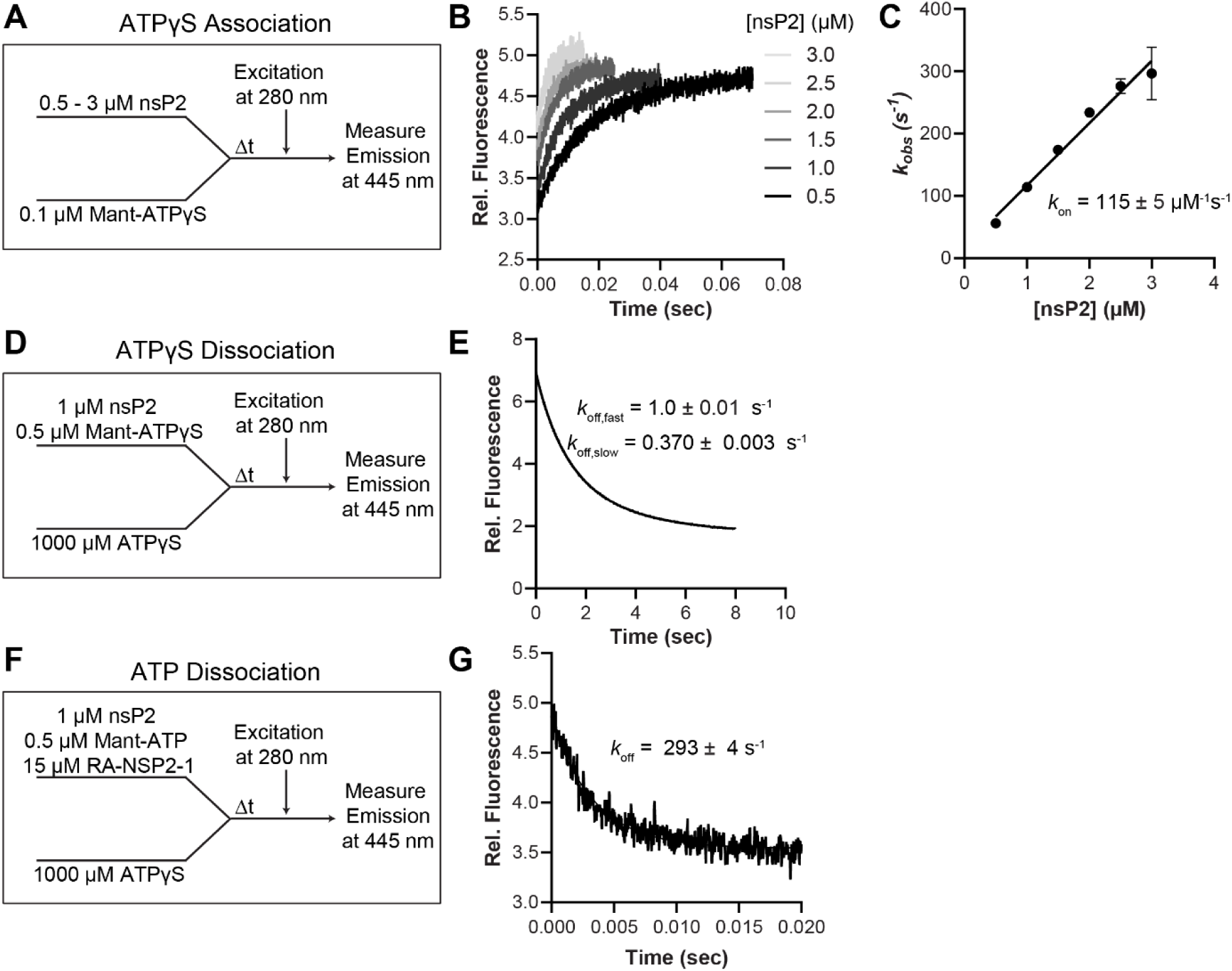
Pre-steady-state kinetics of APT and ATPγS binding to nsP2. **(A)** Exp rimental design for ATPγS association kinetics. nsP2 was rapidly mixed with mant-ATPγS under stopped-flow conditions, and binding was monitored by tryptophan-to-mant FRET. **(B)** ATPγS association kinetics. Representative fluorescence time courses following rapid mixing of mant-ATPγS (0.1 μM) with increasing concentrations of nsP2 (0.5-3 μM). **(C)** Observed rate constants (*k*_obs_) extracted from single-phase fits to the association traces in panel B were replotted as a function of nsP2 concentration (n = 3 independent experiments). Linear regression was used to determine the second-order association rate constant (*k*ₒₙ). **(D)** Experimental design for ATPγS dissociation kinetics. Pre-formed nsP2·mant-ATPγS complexes were rapidly mixed with excess unlabeled ATPγS to initiatw ligand displacement. **(E)** ATPγS dissociation kinetics. Time-dependent loss of sensitized Mant fluorescence following competition with unlabeled ATPγS. Traces were fit to a two-phase exponential decay, revealing fast and slow dissociation components (*k*_off,fast_ and *k*_off,slow_). **(F)** Experimental design for ATP dissociation in the presence of inhibitor. Pre-formed nsP2·mant-ATP complexes were rapidly mixed with excess unlabeled ATP in the presence of the nsP2 inhibitor RA-NSP2- (5 μM). **(G)** ATP dissociation kinetics in the presence of inhibitor. Representative fluorescence decay trace fit to a single-phase exponential model, yielding the apparent ATP dissociation rate constant (*k*_off_).

**Table 3.**
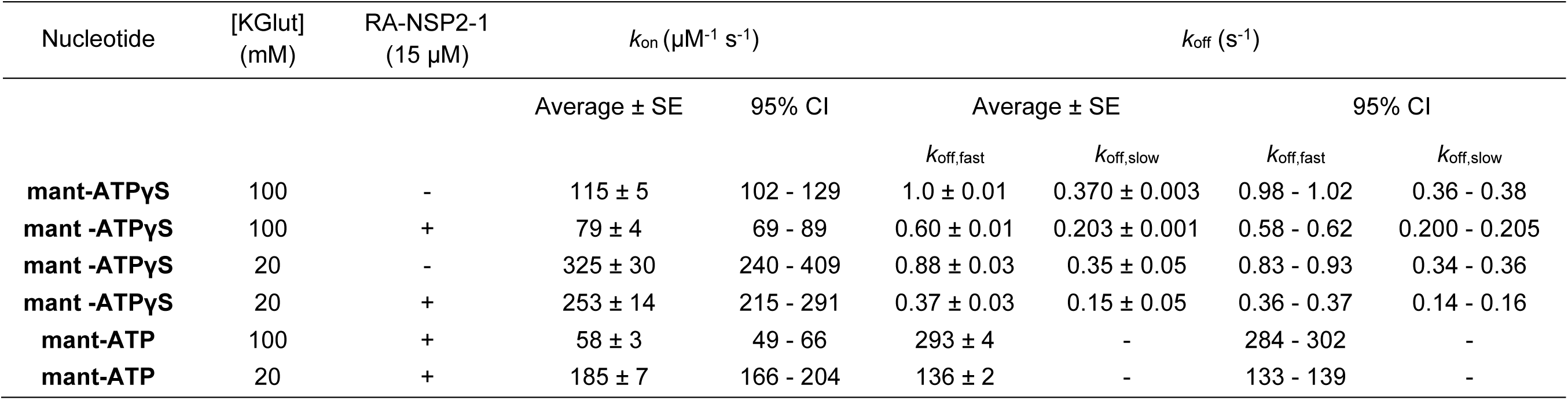
Kinetic parameters for Mant-ATPγ and Mant-ATP to nsP2. Association (*k*_on_) and dissociation (*k*_off_) rate constants for mant-ATPγS and mant-ATP binding to nsP2 were determined by stopped-flow fluorescence under pseudo-first-order conditions. Association kinetics were measured by rapid mixing of mant-ATPγS or mant-ATP (0.1 µM) with nsP2 over a concentration range of 0.5-3 µM, as indicated. Dissociation kinetics were measured by rapid mixing of pre-formed nsP2·mant-nucleotide complexes with execss unlabeled ATPγS (1 mM) to prevent rebinding. Measurements were performed at 20 mM or 100 mM potassium glutamate, as indicated. mant-ATPγS binding kinetics were measured in the absence or presence of 15 µM RA-NSP2-1, whereas mant-ATP kinetics were measured only in the presence of inhibitor to suppress nucleotide hydrolysis. Values are reported as mean ± standard error.

To determine the rate constant for dissociation (*k*_off_) of mant-ATPγS from nsP2, we assembled a nsP2•mant-ATPγS complex and rapidly mixed this complex with sufficient ATPγS to prevent rebinding of mant-ATPγS to nsP2 (**Fig. 4D**). We monitored the loss of fluorescence over time at the excitation and emission wavelengths used above (**Fig. 4D**). The data best fits to a double exponential with rate constants of 1.00 ± 0.01 s^-1^ and 0.370 ± 0.003 s^-1^ (**Fig. 4E**).

The recent discovery of an allosteric inhibitor of nsP2, referred to here as RA-NSP2-1, opened the possibility of monitoring the association and dissociation of mant-ATP (41,42). Furthermore, based on our understanding of the noncompetitive mechanism of action of RA-NSP2-1 (41), ATP binding should not be impacted. The presence of the inhibitor had less than a two-fold effect on the rate constants for association and dissociation at the two ionic strengths evaluated (**Table 3**). The rate constant for mant-ATP association was 58 ± 3 µM^-1^ s^-1^ at 100 mM potassium glutamate and 185 ± 7 µM^-1^ s^-1^ at 20 mM potassium glutamate, both of which were very similar to mant-ATPγS (**Table 3**). Interestingly, the rate constant for mant-ATP dissociation was substantially higher than observed for mant-ATPγS (ompar **Fig. 4E** to **Figs. 4F,G**). The values were: 293 ± 4 s^-1^ and 136 ± 2 s^-1^ at 100 mM and 20 mM potassium glutamate, respectively (**Table 3**).

### Isotope Trapping

The fact that an inhibitor interferes with a step after substrate binding is consistent with data from other helicases in which a conformational change limits the rate of nucleotide hydrolysis (5,37). Given the single-digit micromolar *K*_d_ for ATP inferred from the kinetics of association and dissociation performed above, we could use an isotope-trapping experiment to formally demonstrate the existence of a rate-limiting, conformational-change step (43).

Consider the mechanism shown in **Fig. 5A**. Under the experimental conditions used for the isotope-trapping experiment, nsP2•ATP is in rapid equilibrium with nsP2 and ATP (**Fig. 5B** and **Table 3**). This allowed us to model the pathway beginning from nsP2•ATP. This complex would isomerize into *nsP2•ATP at a rate of *k*_+2_ and return to the ground state at a rate of *k*_-2_. *nsP2•ATP would the n go on to form the nsPs•ADP•Pi product complex at a rate of *k*_+3_. Initiating the reaction by rapidly mixing nsP2 and ATP and quenching the reaction in acid (pulse-quench) will only reveal products that reach nsP2 • DP•Pi or steps thereafter. Over the same time period, chasing the reaction with excess ATP followed by an acid quench (pulse-chase-quench) will reveal both *nsP2•ATP and nsP2•ADP•Pi as the has offers sufficient time to convert *nsP2•ATP to product. Consistent with the existence of a rate-limiting, conformational-change step, more product forms in the pulse-chase-quench experiment than in the pulse-quench experiment (**Fig. 5C**). The solid lines show the time courses predicted by kinetic simulation of the mechanism shown in **Fig. 5A** with the following rate constants: *k*_+2_, 240 s^-1^; *k*_-2_, 30 s^-1^; and *k*_+3_, 40 s^-1^.

**Figure 5.**
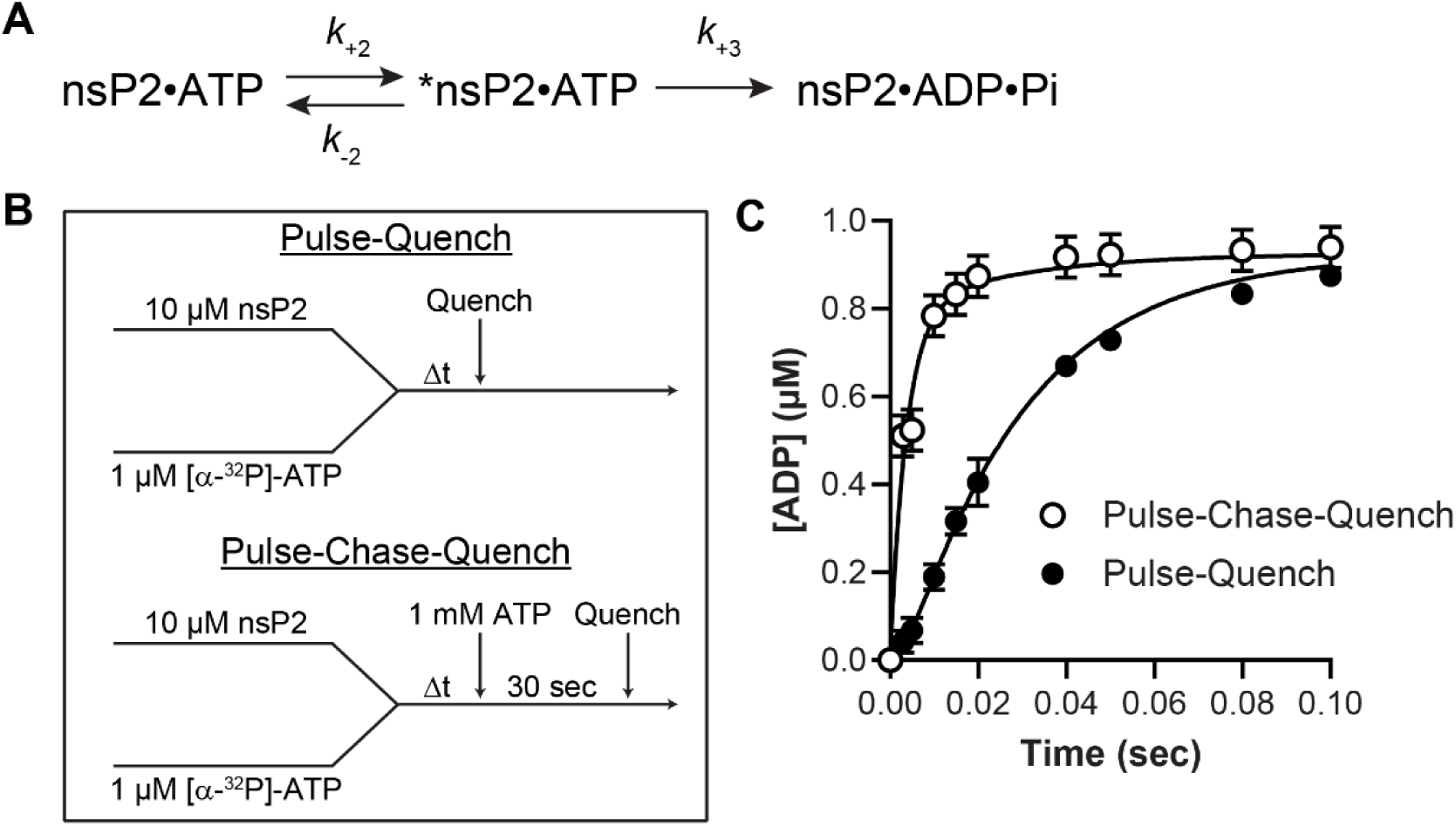
Identification of an isomerized nsP2·ATP complex by isotope trapping. **(A)** Minimal kinetic mechanism for nsP2-catalyzed ATP hydrolysis highlighting the conformational-change and chemical steps defined by rate constants *k*_+2_, *k*_−2_, and *k*_+3_. Pulse-quench and pulse-chase-quench experiments were used to detect formation of an isomerized internal complex (*nsP2·ATP). During the pulse, radiolabeled ATP equilibrates between free nsP2·ATP and the internal *nsP2·ATP complex. Addition of excess unlabeled ATP during the chase removes free nsP2·ATP while trapping isotope in the internal complex, which can proceed to product formation. **(B)** Experimental design for isotope-trapping experiments. In pulse-quench experiments, 10 µM nsP2 was rapidly mixed with [α-³²P]-ATP (1 µM) and quenched with HCl at the indicated time points. In pulse-chase-quench experiments, the nsP2-[α-³²P]-ATP mixture was rapidly chased with excess unlabeled ATP (1 mM), incubated for an additional 30 s, and then quenched with HCl. (C) Product formation and kinetic simulations for pulse-quench and pulse-chase-quench experiments. The additional product observed in pulse-chase-quench experiments provides evidence for formation of the isomerized *nsP2·ATP complex. Solid lines represent global fits of the data to the minimal kinetic mechanism using kinetic modeling, yielding rate constants of *k*_+2_ = 120 s^-1^, *k*_-2_ = 10 s^-1^, and *k*_+3_ = 50 s^-1^. Simulated outputs correspond to formation of nsP2·ADP·Pi (pulse-quench) or total labeled intermediates (*nsP2·ATP + nsP2·ADP·Pi) (pulse-chase-quench).

Our studies of the enterovirus RNA-dependent RNA polymerase have shown a rate-limiting, conformational-change step for nucleotidyl transfer (44). The importance of the quench used to reveal the conformational change became apparent (44). Use of EDTA is essentially like a chase as the EDTA does not disrupt the enzyme complex and provides sufficient time for the intermediate to be converted to product (44). This same complication of using an EDTA quench applies to studies of nsP2. Reactions quenched with EDTA superimpose with the line simulating the pulse-chase-quench experiment (**Fig. 5D**).

### Minimal Mechanism for nsP2-catalyzed Hydrolysis of ATP

We have synthesized the equilibrium, steady-state, and pre-steady-state data into one coherent minimal kinetic mechanism for nsP2-catalyzed hydrolysis of ATP (**Fig. 6**). ATP binds to nsP2 (step 1) to form a complex that isomerizes (step 2). Product is formed in a reaction that is irreversible (step 3). Product dissociation is indicated without any specific order (step 4). However, if Pi binds to the enzyme with the same kinetics of ATP, then Pi would dissociate after ADP at the indicated rate, which is in a time regime that is too fast to affect steady-state turnover. Given the magnitude of the rate constant for the *chemical* step (step 3), it is likely that this step reflects a second conformational-change step that is required for catalysis to occur.

**Figure 6.**
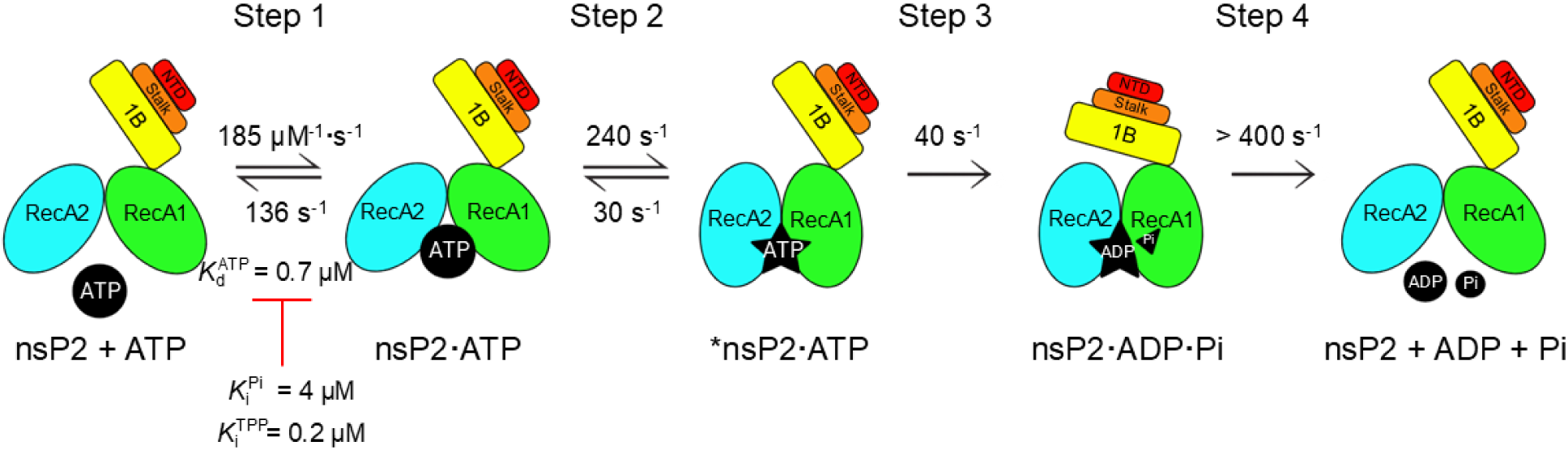
Minimal kinetic mechanism for nsP2-catalyzed ATP hydrolysis. ATP binds to fr nsP2 (Step 1) to form the nsP2•ATP complex, with association and dissociation rate constants of 185 µM⁻¹·s⁻¹ and 136 s⁻¹, respectively (*K*_d_ ≈ 0.74 µM). The complex undergoes a conformational change (Step 2) to form a catalytically omp t nt *nsP2•ATP intermediate (*k*_+2_ = 240 s⁻¹; *k*_−2_ = 30 s⁻¹). Chemistry (Step 3) pro ds irr v rsibly to g n rat nsP2•ADP•Pi (*k*_+3_ = 40 s⁻¹). Product release (Step 4) occurs without an obligatory order. Competitive inhibition constants are indicated for Pi (*K*_i_ ≈ 4 µM) and tripolyphosphat (TPP; *K*_i_ ≈ 0. µM). Rate constants for Step were determined by stopped-flow binding experiments (Fig. 4), those for Steps 2-3 by isotope-trapping analysis (Fig. 5), and product release characteristics by steady-state inhibition studies (Fig. 2). The helicase domain organization of nsP2 is as follows: NTD (residues 1-77), stalk (78-110), 1B (111-175), RecA1 (176-309), and RecA2 (310-440).

### Structural Basis of the Conformational-change Steps Observed Kinetically

Hydrogen deuterium exchange mass spectrometry (HDXMS) studies reported by Sears et al. showed that residues within subdomain 1B (121-132; 151-165), the Walker A motif of the RecA1 domain (250-255), and several other conserved structural elements undergo conformational changes upon nucleotide binding (**Table 4**) (45). In the presence of inhibitor, however, the conformational changes of both 1B and the Walker A motif are selectively suppressed (**Table 4**) (45).

**Table 4.** Assessing comparative protein dynamics by HDXMS and RMSF analysis. Differential HDXMS data, adapted from Sears et al. (45), indicate changes in the dynamics of the specified motifs upon binding ATP or RA-NSP2-1. Comparative protein dynamics were assessed using RMSF analysis of MD simulations to enable direct comparison with the experimental HDXMS data.

| Segment; Residues | HDXMS* |  |  |  | aMD Simulation (RMSF)<sup>†</sup> |  | cMD Simulation (RMSF)<sup>†</sup> |  |
| --- | --- | --- | --- | --- | --- | --- | --- | --- |
|  | nsP2·ATPyS relative to nsP2 |  | nsP2·ATPyS·RA-NSP2-1 relative to nsP2·ATPyS |  | nsP2·ATP relative to nsP2 | nsP2·ATP·RA-NSP2-1 relative to nsP2·ATP | nsP2·ATP relative to nsP2 | nsP2·ATP·RA-NSP2-1 relative to nsP2·ATP |
|  | 1 min | 10 min | 1 min | 10 min |  |  |  |  |
| 1B-1; 121-132 | + | ns | + | ns | ++ | -- | ++ | -- |
| 1B-2; 151-165 | ns | + | ns | ns | ++ | -- | ++ | -- |
| Motif I (Walker A); 182-201 | - | - | - | ns | - | -- | -- | - |
| Motif II (Walker B); 250-255 | ns | - | ns | - | + | - | - | + |
| Motif IIIa; 303-315 | - | - | ns | - | + (dual behavior<sup>‡</sup>) | - | - (dual behavior<sup>‡</sup>) | - (dual behavior<sup>‡</sup>) |
| Motif V; 378-391 | - | ns | - | ns | ++ | -- | -- | - |

\* “+” and “-” indicate a statistically significant increase or decrease in deuterium uptake relative to the reference condition at the specified timepoint (1 min or 10 min); “ns” indicates no significant difference. † ++: $\Delta\text{RMSF} > 0.25 \text{ \AA}$ ; +: $0 < \Delta\text{RMSF} \leq 0.25 \text{ \AA}$ ; -: $-0.25 \text{ \AA} \leq \Delta\text{RMSF} < 0$ ; --: $\Delta\text{RMSF} < -0.25 \text{ \AA}$ . ‡ Upon ATP binding, some residues exhibit both increased and decreased dynamics (dual b

To place these observations in a structural and dynamic framework, we performed both accelerated (aMD) and classical (cMD) molecular dynamics simulations of nsP2 and its complexes. Accelerated MD enhances conformational sampling by lowering energy barriers and raising energetic minima, whereas cMD constrains sampling to the underlying energy landscape, providing a necessary control against overinterpretation of aMD-derived motions (46).

Comparison of the HDXMS data with simulation outputs revealed a high degree of concordance (**Table 4**). The per-residue root mean square fluctuation (RMSF) profiles obtained from aMD (**Fig. 7A**) and cMD (**Fig. 7B**) were nearly identical, supporting the validity of the aMD-derived dynamics. Importantly, aMD more clearly resolved ligand-dependent changes in the magnitude of fluctuations. Subdomains 1B1 and 1B2 exhibited increased dynamics in response to ATP binding (**Fig. 7A**; **Table 4**), consistent with HDXMS. In contrast, the Walker A motif became more rigid upon ATP binding, again in agreement with HDXMS. The addition of the RA-NSP2-1 inhibitor markedly dampened fluctuations in both 1B subdomains and the Walker A motif (**Fig. 7A**; **Table 4**), recapitulating the solution behavior. Thus, ligand-dependent fluctuations observed by aMD align closely with the conformational dynamics measured experimentally.

**Figure 7.**
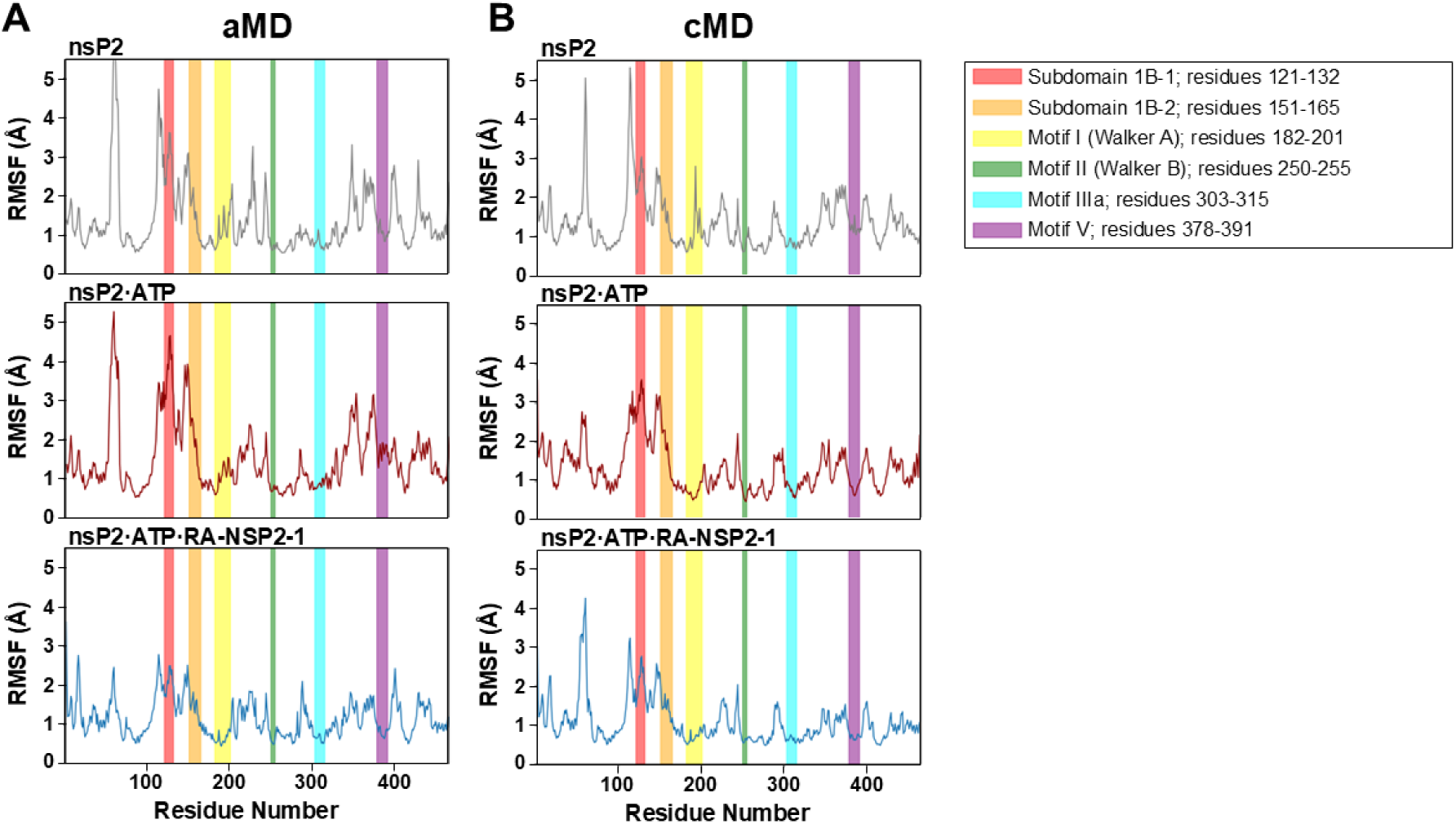
Root mean square fluctuation (RMSF) plots. Cα MSF values for the helicase domain were calculated from (A) accelerated molecular dynamics (aMD) simulations and (B) classical molecular dynamics (cMD) simulations. Colored vertical boxes indicate regions that were differentially deuterated in HDXMS experiments. Residue ranges for the HDXMS segments are as follows: segment 1B-1; 121-132, segment 1B-2; 151-165, motif I (Walker A); 182-201, motif II (Walker B); 250-255, motif IIIa; 303-315, and motif V; 378-391.

A key advantage of MD is the ability to resolve correlated motions between domains. Principal component analysis (PCA) of the aMD trajectories, visualized as porcupine plots (**Fig. 8A**), reveals both the direction and magnitude of these collective motions. In the absence of ligand, the RecA2 domain moves toward RecA1, while 1B moves toward RecA2. ATP binding amplifies these motions, promoting physical interaction between 1B and the domain (nsP2•ATP; **Fig. 8A**). In contrast, inhibitor binding reverses or attenuates these motions, disrupting productive interactions between RecA1, RecA2, and 1B (nsP2•ATP• -NSP2-1; **Fig. 8A**). These relationships are readily apparent in the trajectory animations (**Supplementary Movie 1**). Notably, the inhibitor rigidifies the RecA1-RecA2 interface and prevents 1B from engaging RecA2. Classical MD supports the same conclusions (**Fig. 8B**; **Supplementary Movie 2**).

**Figure 8.**
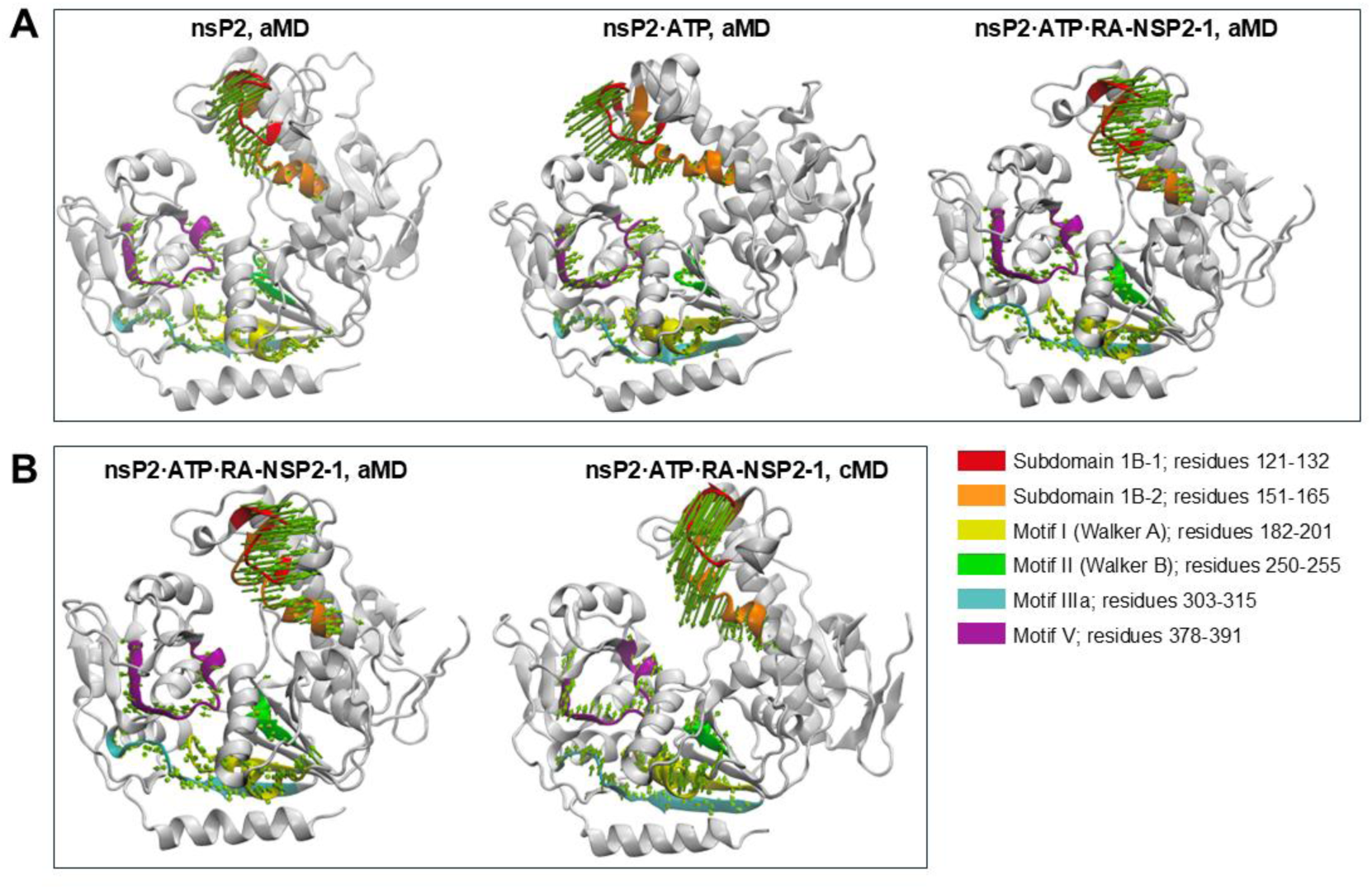
Principal component analysis of nsP2 dynamics from molecular dynamics simulations. (A) Porcupine plots representing the dominant collective motions (principal component 1, PC1) derived from accelerated molecular dynamics (aMD) simulations of nsP2 in different ligand states. Structures are shown for nsP2 (left), nsP2·ATP (middle), and the ternary complex nsP2·ATP·RA-NSP2-1 (right). Vectors (green arrows) indicate the direction and magnitude of backbone atomic displacements corresponding to the principal mode of motion. Structural elements corresponding to subdomain 1B and motifs are highlighted as follows: subdomain 1B segment 1 (red; residues 121-132), subdomain 1B segment 2 (orange; residues 151-165), motif I (Walker A; yellow; residues 182-201), motif II (Walker B; green; residues 250-255), motif IIIa (cyan; residues 303-315), and motif V (purple; residues 378-391). Protein backbone is shown in gray cartoon representation. (B) Comparison of dominant motions for the ternary complex obtained from accelerated molecular dynamics (aMD; left) and classical molecular dynamics (cMD; right) simulations. PC1 vectors are displayed as in panel A, illustrating differences in the magnitude and directionality of correlated motions between enhanced and conventional sampling approaches.

Taken together with the biochemical data showing that the inhibitor blocks the conformational transition required for catalysis, these results point to a specific mechanistic conclusion: catalytic competence requires formation of the 1B-RecA2 interaction. Nucleotide binding promotes this state by driving RecA2 toward RecA1, whereas inhibitor binding prevents it.

### Equilibrium Binding of RNA

The RNA substrates used in this study were developed by our laboratory to study unwinding by the hepatitis C virus (HCV) helicase (4) and later modified slightly to study the West Nile virus (WNV) and Zika virus (ZV) helicases (47). The sequences of the RNAs are provided in **Fig. 9A**. We refer to the strands of the RNA duplex as follows: The strand to which the helicase binds is referred to as the *loading* strand (e.g. U20-ds9-U10 or U40-ds9-U10 in **Fig. 9B**). Because nsP2translocates in the 5’→3’ direction, this enzyme will load on the 5’ side of the duplex. The length of that portion of the loading strand is either 20-nt (U20 in **Fig. 9B**) or 40-nt (U40 in **Fig. 9B**). The other strand is referred to as the *displaced* strand (e.g. ds9-U10 in **Fig. 9B**). An unwinding substrate is formed by annealing the loading and displaced strands (**Fig. 9B**). The “ds9” designation reflects formation of a 9-bp duplex in the unwinding substrate (**Fig. 9B**). The helicase binds to the loading strand, translocate through the duplex, and liberate the displaced strand, which is labeled with fluorescein or ^32^P for binding and unwinding experiments, respectively. To prevent reannealing of the loading and displaced strands, an RNA *trap* strand is added that is identical in sequence to the duplex-forming region of the displaced strand (**Figs. 9A,B**). The goal of these experiments was to determine the equilibrium dissociation constants for these RNAs.

**Figure 9.**
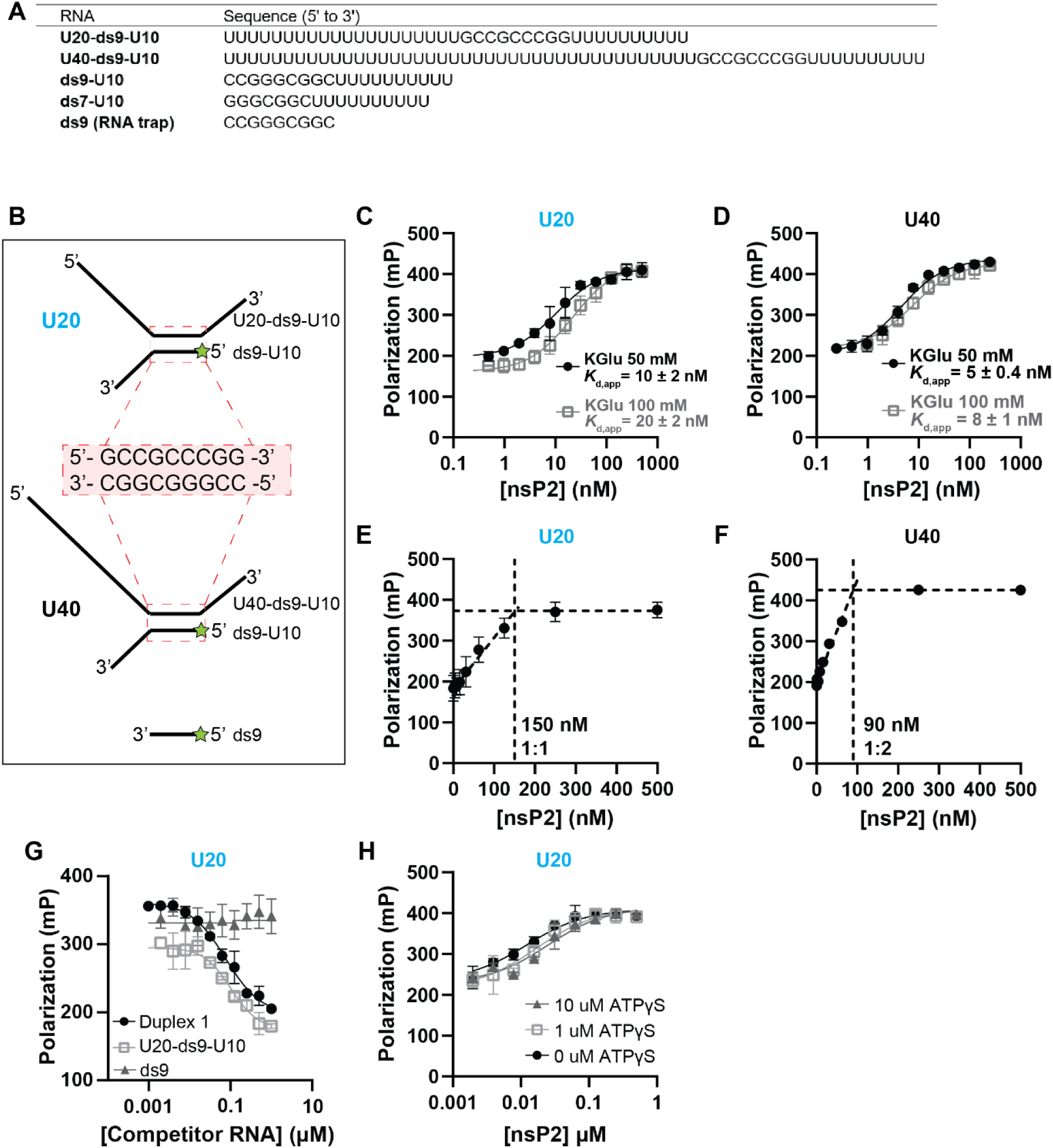
RNA binding by CHIKV nsP2. **(A)** RNA sequences used in fluorescence polarization binding assays and in RNA unwinding assays in Fig. 10. **(B)** Schematic representation of duplex RNA substrates used to assess nsP2 RNA binding. Duplexes contain a 9-bp double-stranded region (ds9; red) flanked by poly(U) single-stranded regions of defined length. The displaced (ds9-U10) strand is 6-FAM-labeled (green star). **(C-D)** nsP2 RNA binding measured by fluorescence polarization. Binding of nsP2 to 1 nM 6-FAM-labeled duplex RNA was measured using U20-ds9-U10/ds9-U10 (C) or U40-ds9-U10/ds9-U10 (D) in either 50 mM or 100 mM potassium glutamate, as indicated. Polarization values were fit to a one-site binding model with offset to obtain apparent dissociation constants (*K*_d,app_). Data are shown as mean ± SD (*n* = 3). **(E-F)** Stoichiometry of nsP2 binding to duplex RNA. nsP2 was titrated into reactions containing U20-ds9-U10/ds9-U10 (E) or U40-ds9-U10/ds9-U10 (F) under conditions where RNA was present at > 10x *K*_d,app_ (150 nM and 90 nM, respectively). Two linear phases were fit by linear regression, and their intersection was used to determine binding stoichiometry. Data are shown as mean ± SD (*n* = 3, E; *n* = 2, F). **(G)** Competitive RNA binding assays. Binding reactions containing 1 nM 6-FAM-labeled U20-ds9-U10/ds9-U10 and 20 nM nsP2 were challenged with increasing concentrations of unlabeled competitor RNA, as indicated. Data are shown as mean ± SD (*n* = 3). **(H)** Effect of ATPγS on nsP2 RNA binding. Fluorescence polarization binding assays with U20-ds9-U10/ds9-U10 were performed in the presence of 0, 1, or 10 µMATPγS.

To measure binding, we used a fluorescence polarization assay (9,48). Binding of nsP2 to fluorescein-labeled RNA caused a substantial change in the magnitude of the fluorescence polarization. We titrated a solution containing either the labeled U20- or U40-containing duplex with nsP2 under equilibrium-binding conditions in which the RNA was present at a concentration on the order of one-tenth the value of the observed dissociation constant. We performed the experiment in the presence of either 50 mM or 100 mM potassium glutamate. We fit the data to a hyperbolic binding equation. The values for the apparent dissociation constants (*K*_d,app_) indicated very tight binding to RNA: 10-20 nM for the U20 duplex (**Fig. 9C**); and 5-8 nM for the U40 duplex (**Fig. 9D**)

The tight binding of RNA to nsP2 provided the opportunity to evaluate binding stoichiometry. In this experiment, RNA was present at a concentration at least 10-fold higher than the observed dissociation constant. Titration of nsP2 under these conditions yielded a stoichiometric-binding isotherm for both duplexes evaluated (**Figs. 9E,F**). We calculated the point of intersection, which revealed a 1:1 RNA:nsP2 complex for the U20 duplex (**Fig. 9E**) and a 1:2 RNA:nsP2 complex for the U40 duplex (**Fig. 9F**). Together, these data are consistent with nsP2 having an RNA-binding-site size of 20-nt.

To determine the extent to which the RNA trap strand will interfere with binding to the unwinding substrate, we titrated the RNA trap strand into a complex of nsP2 and labeled U20 duplex. The RNA trap strand had absolutely no impact on nsP2 binding to the labeled U20 duplex (**Fig. 9G**). As controls, we titrated in the U20 duplex or the U20 loading strand. Both interfered with nsP2 binding to the labeled U20 (**Fig. 9G**).

In our studies of the HCV helicase, we observed a dramatic reduction in the RNA binding affinity of this helicase in the presence of ATP (4). To determine if the affinity of nsP2 for RNA was impacted by the presence of ATP, we performed binding experiments in the presence of a range of concentrations of ATPγS. The effect of nucleotide on RNA binding was minimal (**Fig. 9H**).

### Unwinding of Duplex RNA

Studies of the Dengue virus helicase revealed a peculiar requirement of a 1:2 Mg^2+^:ATP ratio for optimal unwinding by this enzyme (47,49). Our studies of WNV and ZV helicases revealed the same requirement for these enzymes (47). nsP2 unwinding activity was also optimal at a 1:2 Mg^2+^:ATP ratio (**Figs. S3A-G**). nsP2 ATPase activity did not exhibit this peculiarity (data not shown). The sensitivity of the ATPase activity to anions like phosphate motivated us to use acetate and glutamate instead of chloride. Although nsP2 hydrolyzes all four nucleotides with equivalent efficiency (**Table 2**), nsP2 could not utilize GTP for unwinding as well as the other nucleotides (**Figs. S3H,I**).

In order to create a system to study the coupling between the ATPase cycle and unwinding, one needs to define the ATPase cycle and have an RNA duplex that can be unwound in a single step. The HCV helicase moves in steps of three base pairs (4,50). Therefore, one step converts a 9-bp duplex (T_m_ = 50 °C) to 6-bp duplex (T_m_ < 32 °C) (**Fig. 10A**), leading to complete unwinding of the RNA duplex (4). However, using the unwinding assay illustrated in **Fig. 10A**, nsP2 failed to completely unwind the substrates used (**Figs. 10B-D**). Interestingly, the magnitude of substrate unwound increased with increasing length of the loading strand (compare the U20 substrate to the U40 substrate in **Figs. 10C,D**). This phenomenon is well established in the helicase literature and has been referred to as functional cooperativity and suggests that a single nsP2 molecule cannot unwind the 9-bp duplex in a single step. (51,52).

**Figure 10.**
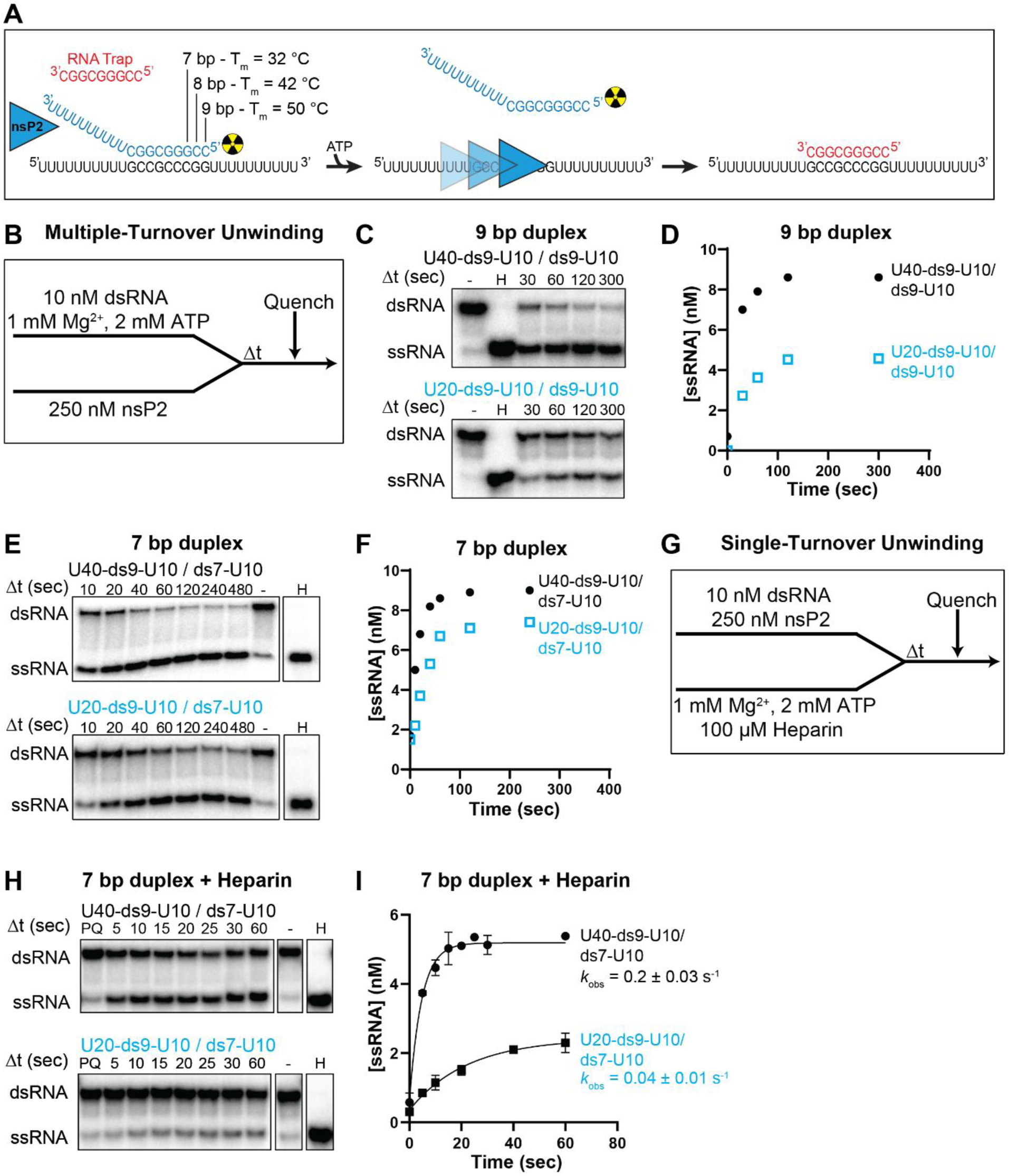
Multiple- and single-turnover RNA unwinding by nsP2. **(A)** Schematic of the nsP2 RNA unwinding assay. Forked duplex substrates were generated by annealing a displayed strand (blue) to a loading strand (black) ontaining ith r a 5′ U 0 or U40 single-stranded region, a 9- or 7-bp duplex-forming r gion, and a 3′ U 0 tail, as defined in Fig. 9A. Annealing with the indicated displaced strand generated duplexes of 9 or 7 bp overall. An RNA trap strand (red) was included to prevent re-annealing of unwound products. RNA sequences are detailed in Fig. 9A. **(B)** Workflow for multiple-turnover RNA unwinding assays. Reactions were performed at 30 °C in the presence of 1 mM Mg²⁺ and 2 mM ATP. **(C)** Representative native gels showing multiple-turnover unwinding of 9-bp duplex substrates (U20-ds9-U10/ds9-U10 and U40-ds9-U10/ds9-U10). **(D)** Quantification of ssRNA product formation from panel C plotted as a function of time. **(E)** Representative native gels showing multiple-turnover unwinding of 7-bp duplex substrates (U20-ds9-U10/ds7-U10 and U40-ds9-U10/ds7-U10). **(F)** Quantification of ssRNA product formation from panel E plotted as a function of time. **(G)** Workflow for single-turnover RNA unwinding assays. nsP2 was pre-associated with duplex RNA prior to rapid mixing with ATP, Mg²⁺, RNA trap, and heparin to initiate unwinding and prevent rebinding of dissociated nsP2. **(H)** Representative native gels showing single-turnover unwinding of 7-bp duplex substrates in the presence of heparin. **(I)** Quantification of single-turnover unwinding kinetics from panel H. Data were fit to a single-exponential model to obtain apparent unwinding rate constants (*k*_obs_). Data represent mean ± SD (*n* = 2).

When we label the displaced strand using [γ-^32^P]-ATP, trace amounts of unconsumed, labeled nucleotide are visible on the native polyacrylamide gel used to separate dsRNA substrate from ssRNA product (**Fig. S4A**). Over time, the labeled ATP is consumed, which means that all the unlabeled nucleotide must have been consumed. So, the inability of nsP2 to unwind all the duplex substrate is likely caused by ATP depletion.

To address the issue of ATP depletion, we pursued the use of ATP regeneration systems. Specifically, we used a system based on a creatine kinase and another based on a proprietary solution purchased from Enzo Life Sciences that we refer to as Enzo (**Fig. S4B**). While they worked well as ATP regeneration systems (**Fig. S4C**), they inhibited RNA unwinding (**Fig. S4D**). It is likely that the phosphate donors of the regeneration systems are binding to nsP2 and competing with ATP as observed for Pi and TPP (**Fig. 6**).

As a last attempt to observe unwinding in a single step by nsP2, we reduced the length of the duplex to 7 bp (**Fig. 10A**). Given the T_m_ value, at 30 °C not all the substrate was in duplex form (see lanes marked by a dash in **Fig. 10E**). However, the duplex was sufficiently stable to assess unwinding efficiency. Under multiple turnover conditions and before all the ATP in the reaction could be consumed, 70-90% of the substrate could be unwound with a much smaller difference observed between the 20-nt and 40-nt loading strands (**Fig. 10F**). We conclude that some fraction of the product formed using the 7-bp duplex occurs in a single binding event, perhaps using a single step.

To study the fraction of the nsP2-RNA complex that turns over in a single binding event, we performed a single-turnover experiment. nsP2 was incubated with the 7-bp duplex substrates and mixed with ATP, Mg^2+^, and heparin (**Fig. 10G**). If heparin is present in the reaction at the time of nsP2 addition, the reaction is completely inhibited (**Fig. S4E**). Therefore, any nsP2 that is not bound to RNA at the start of the reaction or dissociates from RNA during the reaction will be inhibited. Using the U20 loading strand, 20% of the substrate was unwound. Using the U40 loading strand, 50% of the substrate was unwound (**Figs. 10H,I**). However, under these conditions, 100% of the RNA should have been bound based on our understanding of nsP2 RNA binding affinity and stoichiometry (**Fig. 9**). It is possible that the alignment of nsP2 on the RNA is heterogeneous. The magnitude of the observed rates of unwinding was 0.04-0.2 s^-1^ (**Fig. 10I**). This range is too slow to be a process that is coupled to ATP hydrolysis. Additional studies will be required to create a substrate whose unwinding rate is coupled directly to the ATPase cycle.

## Discussion

Alphaviruses represent a continuing and expanding threat to public health, yet mechanistic understanding of alphavirus-encoded enzymes remains limited relative to other positive-strand RNA viruses. One consequence of this gap is the absence of robust biochemical frameworks that can be used to interrogate enzyme mechanism or guide antiviral discovery. Here, we provide such a framework for the alphavirus nsP2 NTPase/helicase by defining the kinetic steps that govern nucleotide hydrolysis and by placing these steps in the context of the structure and dynamics of this enzyme (**Fig. 6**).

A central observation is that nsP2 binds nucleoside triphosphates with unusually high affinity. This property distinguishes nsP2 from many helicases and enabled single-turnover, isotope-trapping, and pre-steady-state experiments that are rarely feasible for viral helicases. The use of mant-nucleotides to quantify binding and kinetics follows directly from the approach pioneered by Webb and colleagues for SF1 helicases, where tryptophan-to-mant fluorescence resonance energy transfer (FRET) has proven uniquely powerful for resolving nucleotide association and dissociation kinetics, as well as equilibrium binding (37). In the context of nsP2, these experiments reveal rapid binding, with association rate constants approaching diffusion limits and dissociation rate constants consistent with low-micromolar affinity. Thus, ATP binding is not rate-limiting for nsP2, and the enzyme spends most of its catalytic cycle in a nucleotide-bound state.

The binding data also provides an unambiguous conclusion regarding the origin of nucleotide affinity. Tripolyphosphate binds nsP2 with affinity comparable to that of ATPγS and far greater than that of inorganic phosphate or DP, and it acts as a competitive inhibitor without serving as a substrate (**Fig. 2G, S2E**). These observations demonstrate that the triphosphate moiety is the primary driver of NTP binding. Consistent with this conclusion, nsP2 hydrolyzes all canonical NTPs with similar efficiency and can support RNA unwinding with multiple NTPs (**Table 2, Fig. S3H,I**). Base-specific interactions, therefore, play a minimal role in substrate recognition. This feature may be advantageous in the replication organelle, where nucleotide pools could fluctuate, but it also creates a potential liability. In principle, a cytoplasmic nsP2 enzyme capable of hydrolyzing all NTPs could deplete cellular nucleotide pools. The strong inhibition by inorganic phosphate suggests a solution to this problem. Cytoplasmic phosphate concentrations are orders of magnitude higher than the measured Ki, implying that Pi would suppress nsP2 activity outside the replication compartment (53). Thus, Pi may serve as a metabolite-based safeguard that prevents unregulated NTP hydrolysis by nsP2 in the cytoplasm, while the specialized environment of the spherule permits productive activity.

The kinetic analysis places these binding properties into a mechanistic framework. ATP binding is followed by a reversible isomerization, revealed by isotope-trapping experiments that demonstrate accumulation of an internal ATP-bound state (**Fig. 5**). This observation aligns with prior studies of SF1B helicases, particularly the RecD2 system studied by Webb and colleagues, in which a conformational change following nucleotide binding precedes hydrolysis (37). Structural studies of SF1B helicases, including RecD2 and coronavirus nsp13, have shown that ATP binding induces rearrangement of the RecA-like motor domains, providing a structural basis for this isomerization (35,36,54). In this regard, nsP2 conforms to the canonical SF1 helicase paradigm.

However, the nsP2 mechanism extends this framework by identifying a second conformational transition that limits catalysis. The step regulating product formation is irreversible (**Fig. 2H,I**) and occurs on the millisecond timescale. Product release is too rapid to account for the observed single-turnover rate (**Fig. 2D-F**). These observations are consistent with the rate-limiting step for ATP hydrolysis reflecting a conformational change required to achieve the catalytically competent state.

The structural dynamics governing the observed conformational changes is informed by the molecular dynamics simulations. ATP binding stabilizes the catalytic core, particularly the Walker A motif, while increasing the dynamics of subdomain 1B in the N-terminal region (**Table 4**, **Fig. 7**). This redistribution of dynamics suggests that ATP binding creates a state in which the active site is ordered while an accessory domain becomes mobile and capable of creating the catalytically competent state. Such behavior is consistent with the broader helicase literature, where accessory domains frequently regulate the activity of the RecA motor core. For example, in Pif1-family helicases, ATP and nucleic acid binding drive large conformational changes in accessory domains that are required for function, although these transitions have not been resolved kinetically to the same extent (55). Similarly, studies of nsp13 emphasize extensive interdomain flexibility and allosteric coupling but do not define discrete kinetic steps (54).

The availability of the nsP2 inhibitor RA-NSP2-1 provides an important tool for interrogating this transition (41,42). This compound has been characterized previously as an allosteric inhibitor of nsP2 and does not interfere substantially with nucleotide binding (41). In our hands, MD simulations suggest that the inhibitor suppresses the ATP-dependent dynamics of subdomain 1B and further stabilizes the nucleotide-bound state of the catalytic core. These observations indicate that the inhibitor traps nsP2 in a pre-catalytic conformation in which ATP is bound and the initial isomerization has occurred, but the subsequent conformational transition required for catalysis is blocked. Thus, RA-NSP2-1 does not reveal the mechanism de novo but serves as a powerful probe that allows assignment of the rate-limiting step to a conformational transition involving the N-terminal/1B region.

The use of small-molecule inhibitors to trap specific conformational states is well established in other systems. For example, inhibitors of the HCV NS3 helicase and SARS-CoV-2 nsp13 have been shown to stabilize distinct conformations of the helicase core or accessory domains, thereby providing insight into the coupling between nucleotide binding and mechanical function (36,49). In these systems, inhibitors have been used to bias conformational equilibria and reveal otherwise transient states (36). The behavior of RA-NSP2-1 is consistent with this paradigm, reinforcing the conclusion that nsP2 activity is governed by conformational selection rather than simple chemical kinetics.

The RNA-binding and unwinding experiments further emphasize that nsP2 function cannot be understood solely in terms of ATP hydrolysis. nsP2 binds RNA with high affinity and a defined binding-site size but unwinding under single-turnover conditions is slow relative to ATP hydrolysis. This result suggests that formation of a productive nsP2•RNA complex is itself a limiting process and that ATP hydrolysis is not tightly coupled to strand separation under the conditions tested. Similar phenomena have been observed for other helicases, where substrate architecture, protein oligomerization, and accessory factors strongly influence unwinding efficiency (5,6). In the case of nsP2, the requirement for longer loading strands and the evidence for functional cooperativity suggest that multiple nsP2 molecules or specific RNA configurations may be required for efficient unwinding (**Fig. 10**).

Taken together, these findings may suggest that nsP2 is not optimized to function as an autonomous helicase operating in solution. Precursor forms of nsP2, complexes of nsP2 with other non-structural proteins, and/or the context of the replication spherule may be required for efficient coupling of nucleotide hydrolysis to RNA unwinding. The tight binding of nucleotide, sensitivity to phosphate, and requirement for a conformational transition involving the N-terminal/1B region all point to a mechanism in which ATP hydrolysis is gated by structural and possibly environmental cues. Such regulation would ensure that energy expenditure is coupled to productive events, such as RNA remodeling or replication-complex function, and would prevent deleterious interactions with cellular RNAs.

In summary, nsP2 hydrolyzes NTPs through a mechanism in which tripolyphosphate-driven binding is followed by two conformational transitions, the second of which limits catalysis and is sensitive to allosteric inhibition. This mechanism is consistent with core features of SF1B helicases while revealing an additional layer of regulation that may be specific to alphavirus biology. These findings provide a foundation for understanding nsP2 function and offer a framework for future studies aimed at linking ATP hydrolysis to RNA unwinding and viral replication.

## Supporting Information

Supporting information is online.

## Data Availability

All data are incorporated into the article and its online supplementary material. Constructs and data sets presented in this study are available upon request.

## Acknowledgements

We would like to acknowledge members of the Cameron-Arnold laboratory for helpful conversations and discussions throughout this study. We also thank the READDI-AC virology team, including John D. Sears, Nathanial J. Moorman, and Mark T. Heise, for sharing their results on nsP2 biology and biophysics prior to publication. Additionally, we thank Kenneth A. Johnson for thoughtful feedback on kinetic modeling.

## Abbreviations

aMD: accelerated molecular dynamics
ATP: adenosine triphosphate
ATPγS: adenosine 5′-O-(3-thiotriphosphate)
CHIKV: Chikungunya virus
cMD: classical molecular dynamics
CTP: cytidine triphosphate
dsRNA: double-stranded RNA
EDTA: ethylenediaminetetraacetic acid
FRET: Forster resonance energy transfer
GTP: guanosine triphosphate
HCV: hepatitis C virus
HDXMS: hydrogen deuterium exchange mass spectrometry
HEPES: 4-(2-hydroxyethyl)-1-piperazineethanesulfonic acid
KCl: potassium chloride
MD: molecular dynamics
NMR: nuclear magnetic resonance
NPT: constant pressure and temperature ensemble
NTP: nucleoside triphosphate
NTPase: nucleoside triphosphate hydrolase
NVT: constant volume and temperature ensemble
PAGE: polyacrylamide gel electrophoresis
PCA: principal component analysis
PEI: polyethylenimine
Pi: inorganic phosphate
PMEMD: Particle Mesh Ewald Molecular Dynamics
RMSF: root mean square fluctuation
RNA: ribonucleic acid
RTPase: RNA triphosphatase
SF1B: superfamily 1B
SUMO: small ubiquitin-like modifier
TBE: Tris-borate-EDTA
TCEP: tris(2-carboxyethyl)phosphine
TLC: thin-layer chromatography
TPP: tripolyphosphate
UTP: uridine triphosphate
VMD: Visual Molecular Dynamics
WNV: West Nile virus
WT: wild type
ZV: Zika virus

---

**Figure S1.**
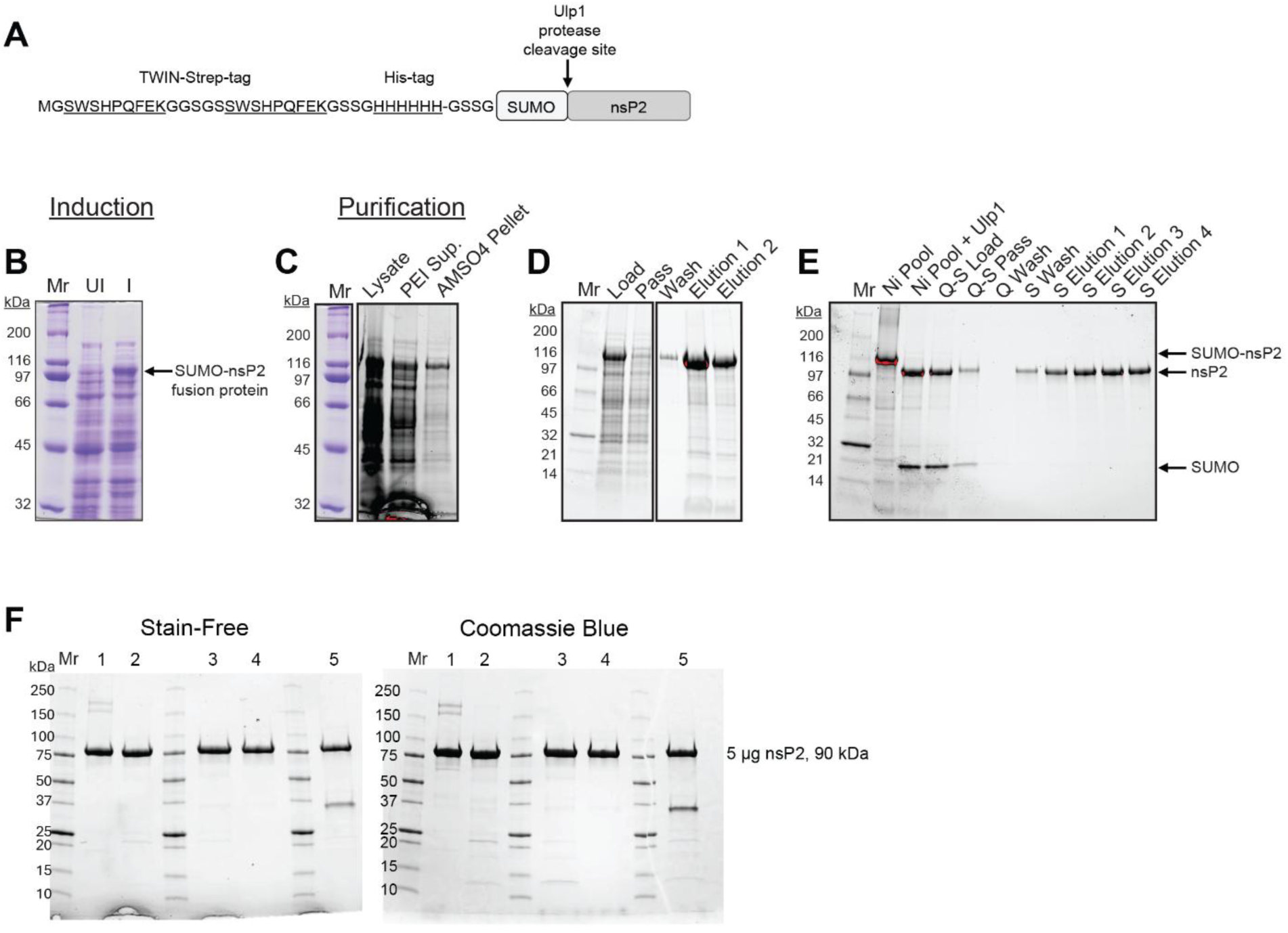
Expression and purification of recombinant SUMO-nsP2. **(A)** Schematic of the Twin-Strep-6xHis-SUMO-CHIKV-nsP2 expression construct used in this study. nsP2 was released from the affinity and solubility tags by Ulp1 SUMO protease cleavage. **(B-E)** Expression and purification workflow. Recombinant protein was expressed in *E. coli* for 48 h at 15 °C prior to cell harvest. Cell pellets were lysed by high-pressure homogenization, followed by treatment with polyethylenimine (PEI) and ammonium sulfate to remove nucleic acids and concentrate protein. Clarified lysates were applied to a Ni²⁺-NTA affinity column, and bound protein was eluted and subjected to Ulp1 digestion to remove the SUMO tag. Cleaved protein was further purified by sequential anion (Q) and cation (S) exchange chromatography to separate nsP2 from the SUMO fragment and contaminants. **(F)** SDS-PAGE analysis of 5 µg purified nsP2 from five independent preparations. Preparations 1-4 were generated using the full purification workflow described above and were used for all biochemical experiments. Preparation 5 utilized Ni²⁺-NTA affinity chromatography followed by size-exclusion chromatography in place of Q and S ion exchange purification and is shown for comparison only.

**Figure S2.**
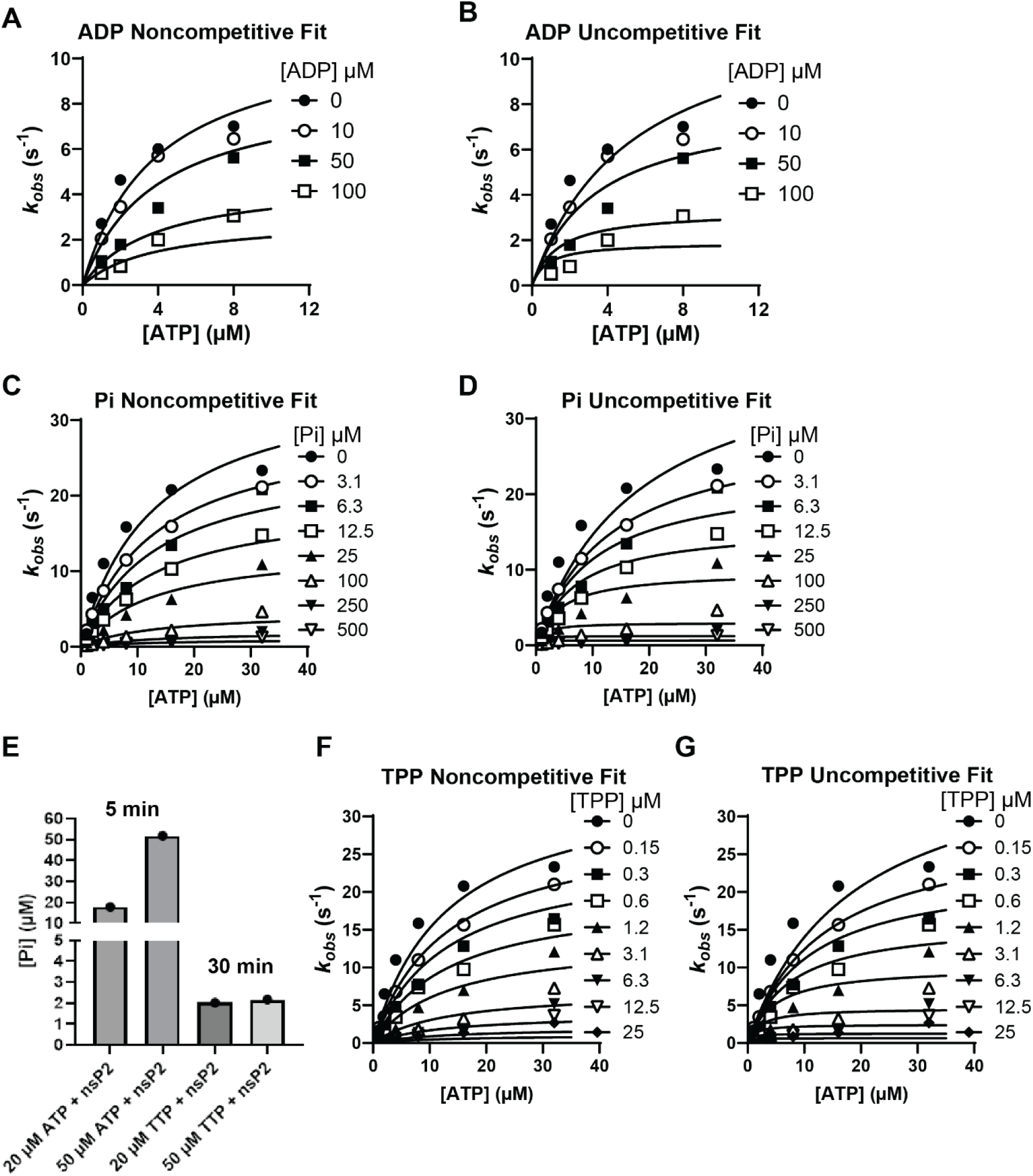
Alternative inhibition models and evaluation of TPP as a substrate. **(A-D, F-G)** Alternative inhibition models for nsP2 ATPase activity. Steady-state ATPase data shown in Fig. 2 (panels D [ADP], E [Pi], and G [TPP]) were converted to observed rate constants (kobs, s⁻¹) by normalizing initial velocities to the enzyme concentration (0.5 nM nsP2) and plotted as a function of ATP concentration. Data were globally fit by nonlinear regression to noncompetitive and uncompetitive inhibition models for comparison with the competitive inhibition models shown in Fig. 2. (A) ADP noncompetitive inhibition model. (B) ADP uncompetitive inhibition model. (C) Pi noncompetitive inhibition model. (D) Pi uncompetitive inhibition model. (F) TPP noncompetitive inhibition model. (G) TPP uncompetitive inhibition model. **(E)** Evaluation of TPP as a substrate for nsP2. Inorganic phosphate (Pi) production was measured using a malachite green colorimetric assay. Free phosphate binds malachite green, producing an increase in absorbance at 620 nm. nsP2 (20 nM) was incubated with ATP or TPP at the indicated concentrations (20 or 50 µM). Reactions were quenched with EDTA at 5 or 30 minutes and mixed with malachite green reagent prior to absorbance measurement at 620 nm. Bars represent Pi produced under each condition. No detectable Pi production was observed with TPP relative to ATP.

**Figure S3.**
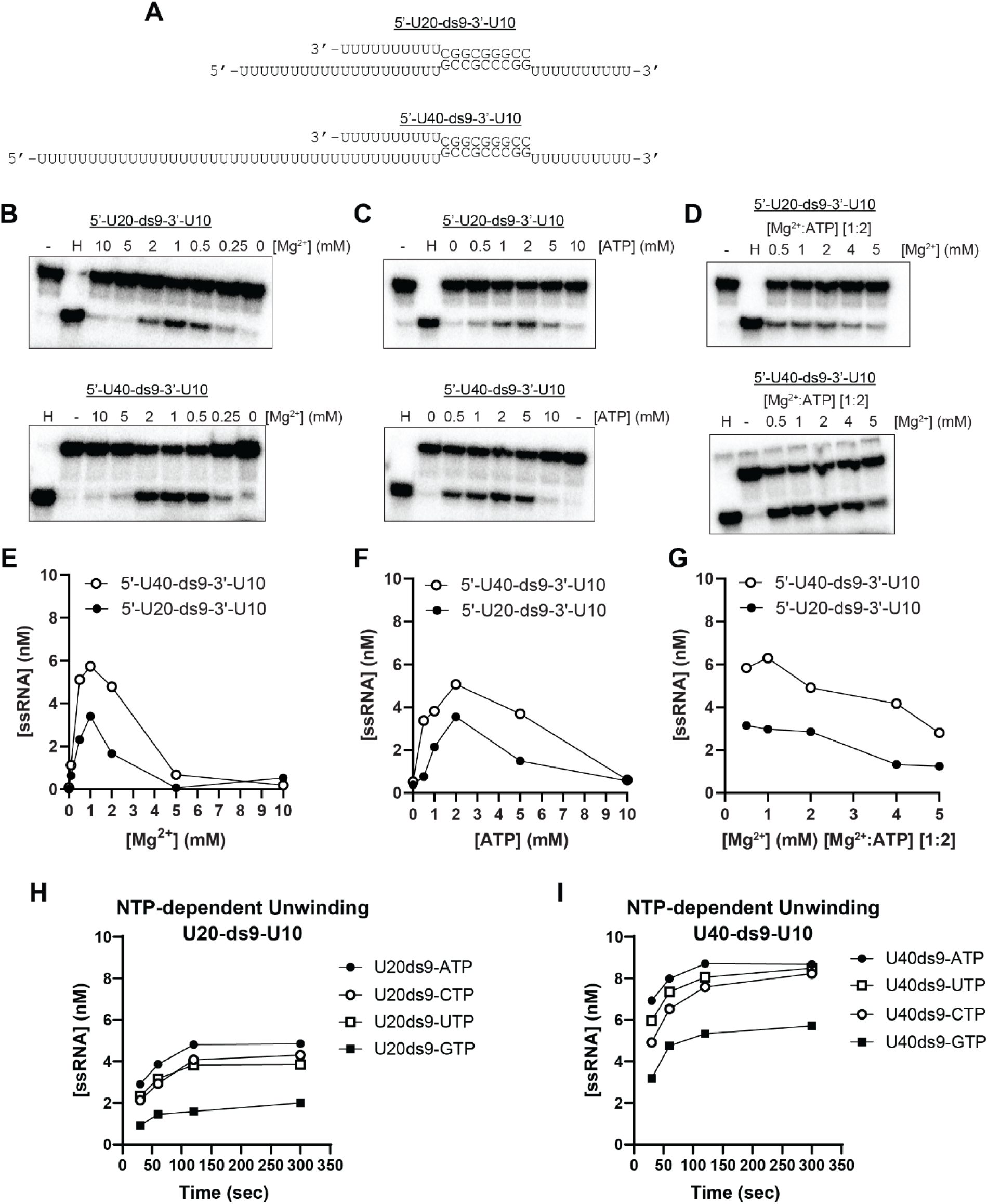
Optimization of nsP2 RNA unwinding assay conditions. **(A)** RNA substrates used for optimization experiments. Forked duplex substrates contained either a U20 or U40 5′ loading strand annealed to a displaced strand to form a 9-bp duplex region with a 3′ U10 tail, as defined in Fig. 9A. **(B, E)** Magnesium dependence of steady-state RNA unwinding with ATP fixed at 2 mM. Representative native gels (B) show unwinding reactions performed in 25 mM HEPES (pH 7.5), 20 mM potassium glutamate, 10 mM TCEP, 0.5 µM RNA trap, 10 nM radiolabeled RNA substrate, and 250 nM nsP2, with magnesium acetate titrated from 1-10 mM. Quantification of ssRNA product formation is shown in panel E. **(C, F)** ATP dependence of steady-state RNA unwinding with magnesium fixed at 1 mM. Representative native gels (C) show reactions performed under the same conditions as in panel B, with ATP titrated from 1-10 mM. Quantification of ssRNA product formation is shown in panel F. **(D, G)** Effect of varying Mg²⁺ and ATP concentrations while maintaining a constant Mg²⁺:ATP ratio of 1:2. Representative gels (D) show reactions performed with magnesium acetate varied from 0.5-5 mM and ATP adjusted accordingly to maintain the 1:2 ratio. Quantification of ssRNA product formation is shown in panel G. **(H-I)** Nucleotide specificity of nsP2-mediated RNA unwinding. Steady-state unwinding assays were performed under optimized conditions using ATP, CTP, GTP, or UTP with U20-ds9-U10 (H) or U40-ds9-U10 (I) substrates.

**Figure S4.**
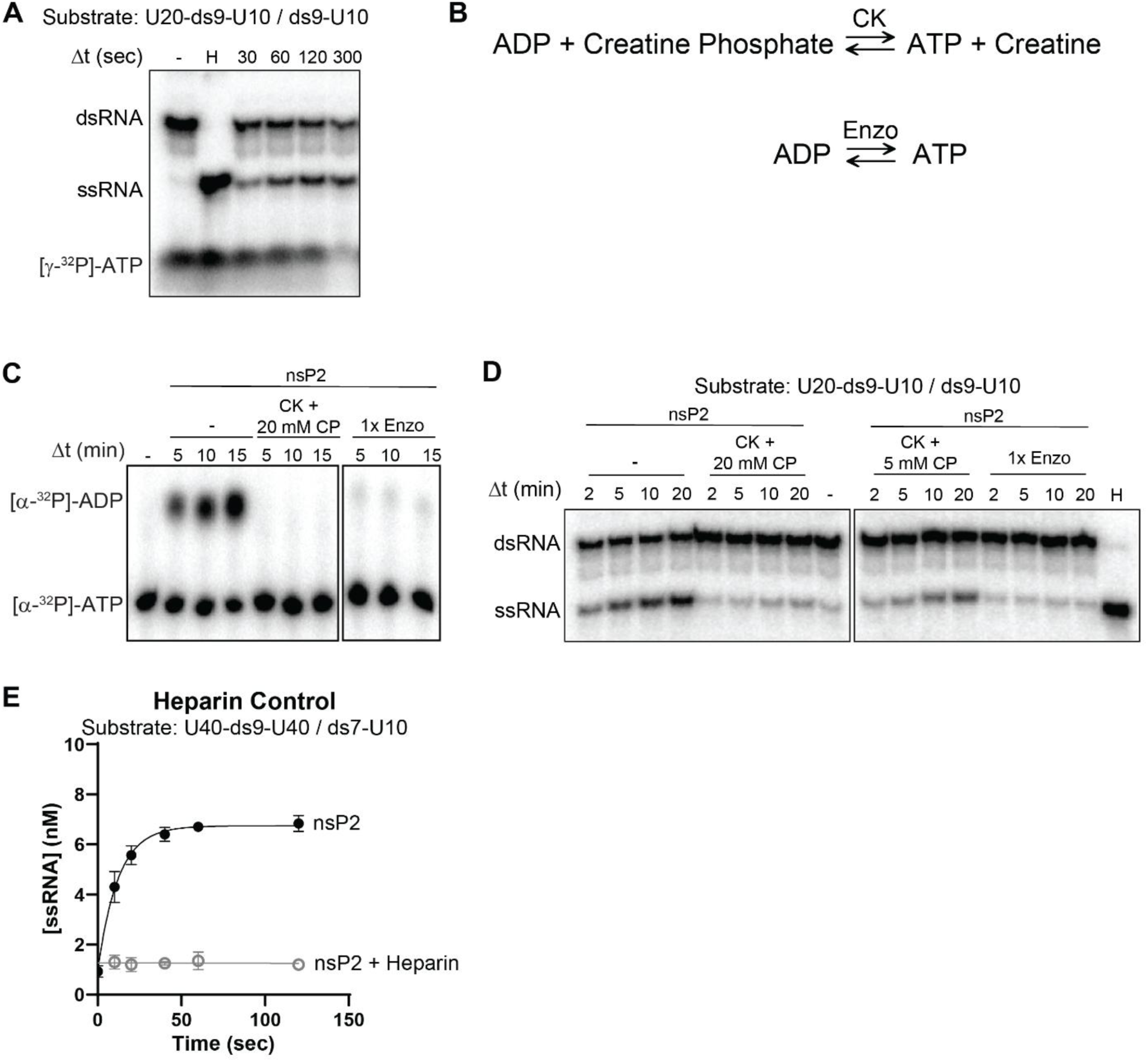
ATP depletion, ATP regeneration systems, and heparin control experiments. **(A)** Full native PAGE gel corresponding to the multiple-turnover unwinding conditions shown in Fig. 10C (U20-ds9-U10/ds9-U10 substrate). In addition to dsRNA and ssRNA species, residual [γ-³²P]-ATP derived from RNA labeling is visible at early time points and progressively depleted between 100-300 s. Depletion of ATP coincides with the plateau observed in multiple-turnover unwinding time courses. **(B)** Schematic representation of ATP regeneration systems used in this study. The creatine kinase (CK) system regenerates ATP from ADP using creatine phosphate (ADP + CP ⇌ ATP + creatine). The proprietary Enzo system regenerates ATP from ADP via an enzyme-coupled reaction. **(C)** Steady-state ATPase assays performed in the presence of ATP regeneration systems, monitored by TLC. Reactions contained [α-³²P]-ATP and nsP2 and were supplemented with either CK + creatine phosphate (CP) or 1x Enzo regeneration system, as indicated. Regeneration systems maintain ATP levels and suppress net accumulation of ADP, demonstrating effective ATP recycling under ATPase assay conditions. **(D)** Effect of ATP regeneration systems on RNA unwinding. Multiple-turnover unwinding assays were performed using the U20-ds9-U10/ds9-U10 substrate in the presence of CK + CP (5 or 20 mM) or 1× Enzo system. Representative native gels are shown. Inclusion of ATP regeneration systems reduces apparent unwinding despite maintaining ATP levels. **(E)** Heparin trap control for single-turnover unwinding assays. Unwinding of the U40-ds9-U40/ds7-U10 substrate was performed in the presence or absence of 100 µM heparin under conditions described in Fig. 10G-I. Heparin suppresses unwinding by trapping free nsP2, confirming enforcement of single-turnover conditions.

